# Genomic signatures of reproductive isolation are decoupled from floral divergence in a long-standing hybrid zone

**DOI:** 10.64898/2026.01.17.700129

**Authors:** Benjamin W. Stone, N. Hoyt Williams, Trinity H. Depatie, Zachary J. Radford, Alice M. Mosley, Carolyn A. Wessinger

## Abstract

A central goal in evolutionary biology is to understand how species boundaries are maintained despite gene flow. Two North American wildflower species with divergent floral syndromes, *Penstemon davidsonii* (bee syndrome) and *P. newberryi* (bird syndrome), have formed hybrid zones in the eastern Sierra Nevada for at least 85 years. We combined multiple approaches to test whether divergent floral syndromes enforce reproductive isolation as predicted by classic models of pollinator-driven ecological speciation. The two parent species exhibited strong divergence across multivariate trait space and have maintained genomic differentiation despite persistent hybridization. Floral hue, a key component of pollination syndrome, mapped to a single genomic region containing two candidate genes with large effects on anthocyanin pigment composition. Pollinator visual models indicated that genetic variation at this region affects detectability to hummingbirds, but not bees. Genomic cline analyses identified many significantly steep clines, suggesting a polygenic basis to reproductive isolation. Surprisingly, these barriers appear unrelated to floral isolation; the major floral hue locus exhibits a strikingly shallow genomic cline and elevated heterozygosity, suggesting pervasive gene flow. Our results reveal that conspicuous trait divergence can be decoupled from reproductive barriers, challenging assumptions about how reproductive isolation is maintained in hybridizing populations.

## Introduction

Local adaptation to divergent ecological conditions can generate barriers to reproduction that are often incomplete (Butlin & Faria, 2024). Such incomplete reproductive isolation can give rise to hybrid zones when genetically distinct species come into contact and produce offspring (Barton & Hewitt, 1985). Often viewed as “natural laboratories” where reproductive isolation between species is incomplete (Hewitt, 1988), the natural recombination of distinct phenotypes and genotypes in hybrid zones can be leveraged to identify the genetic basis of trait variation and reproductive barriers. Hybrid zones have increasingly been used to reveal the genomic architecture of reproductive isolation and local adaptation (e.g., Powell et al., 2021; Schield et al., 2024; Semenov et al., 2025), to identify the traits and ecological gradients that generate or reinforce barriers to gene flow (Sianta et al., 2024; e.g., Stubbs et al., 2023), and to map the genetic basis of phenotypic divergence itself (e.g., Hooper et al., 2024; Justen et al., 2024; Schield et al., 2024). Hybrid zones thus provide a unique lens through which the evolutionary processes that maintain or erode species boundaries may be studied.

Hybrid zones are typically considered ephemeral, eventually leading to either the collapse of species barriers through extensive hybridization and introgression (fusion) or the completion of speciation through selection against unfit hybrids (reinforcement). In some cases, hybrid zones experience neither fusion nor reinforcement over long periods of time (e.g., Aguillon & Rohwer, 2022; DeRaad et al., 2025; Dougherty & Carling, 2024; Semenov et al., 2025). These persistent, long-standing hybrid zones are of interest because they represent stable equilibria in which species boundaries have not collapsed, yet reproductive isolation is also incomplete. Long-standing hybrid zones may be tension zones, which are maintained through a balance of parent species dispersal and selection against hybrids (Barton & Hewitt, 1985; Key, 1968; Slatkin, 1973). Alternatively, they may be dispersal-independent, and maintained, for example, through “bounded hybrid superiority”, in which hybrids can outcompete either parent species in narrow environments where competition is weak (Moore, 1977). The genetic, ecological, and evolutionary mechanisms underlying why some hybrid zones persist in this state of balance, seemingly suspended in time, while others do not, remains an enduring mystery of evolutionary biology.

One of the best-documented long-standing hybrid zones is formed between two western North American wildflower species, *Penstemon davidsonii* Greene and *P. newberryi* A.Gray (Clausen et al., 1940; García et al., 2023; Kimball, 2008; Kimball et al., 2008; Kimball & Campbell, 2009). As members of *Penstemon* subgenus *Dasanthera* (the “shrubby beardtongues”), these species are well-known for their interfertility and propensity to hybridize when in sympatry with close relatives (Stone et al., 2023; Stone & Wessinger, 2024; Stone & Wolfe, 2021). The distributions of these species overlap in the Sierra Nevada and the southern Cascade Range, where *P. davidsonii* is found only at very high elevations, typically above the treeline, and *P. newberryi* is found at lower elevations. These species have diverged across a constellation of traits, including physiology (Kimball & Campbell, 2009), growth habit (Clausen et al., 1940), and perhaps most notably, floral traits typically associated with distinct pollinator communities (García et al., 2023; Kimball, 2008; Kimball et al., 2008). In particular, *P. newberryi*, with its narrow, pink-magenta flowers and exserted stamens and styles, represents a typical hummingbird-pollinated syndrome, while *P. davidsonii*, with its wide, blue-purple flowers and inserted stamens and styles, represents a typical hymenopteran-pollinated syndrome (Kimball, 2008). In many angiosperm lineages, these pollination syndromes – complex multi-trait adaptations to specific types of pollinators – have been assumed to generate reproductive isolation ("floral isolation") through a model of pollinator-driven ecological speciation (Van Der Niet et al., 2014). Under this model, local adaptation to distinct pollinator communities leads to assortative mating that confers reproductive isolation in secondary contact. Yet, since at least the early 20th century, *P. davidsonii* and *P. newberryi* have repeatedly formed natural hybrid zones without any evidence of intrinsic reproductive incompatibilities (Clausen et al., 1940). More recent work in this system found that F1 and later generation hybrids have similar seed sets as parent species, further supporting the lack of intrinsic reproductive barriers to hybridization (Kimball et al., 2008), and that F1 hybrids also have intermediate physiological traits related to elevational adaptation (such as water use efficiency and maximum photosynthetic rate), which may explain their persistence at intermediate elevations where they are found (Kimball & Campbell, 2009).

Species boundaries may be maintained even in the absence of intrinsic reproductive incompatibilities. Even if hybrids are fertile, the divergent pollination syndromes in *P. davidsonii* and *P. newberryi* could maintain species boundaries if hybrids experience reduced visitation due to pollinator discrimination against mismatched combinations of floral traits. Prior work identified only partial overlap in the visiting pollinator communities, leading to the hypothesis that floral traits may shape patterns of gene flow in this system (Kimball, 2008). In addition to visual cues, floral scent divergence may also drive differences in pollinator visitation and thus patterns of gene flow; although *P. davidsonii* and *P. newberryi* produce similar total emissions, they do differ in certain characteristics of scent, including scent composition and aromatic emission (García et al., 2023). Notably, hummingbirds visit *P. newberryi* and hybrids, but not *P. davidsonii* (Kimball, 2008). While the abundance of *P. newberryi*-like phenotypes in hybrids suggests that gene flow may be elevated between hybrids and *P. newberryi* compared to the other backcross direction (Kimball, 2008), to date, no genetic data have been used to test this hypothesis.

The *P. davidsonii*–*P. newberryi* hybrid zone exemplifies the conundrum of long-standing hybrid zones: why do species boundaries persist when barriers to gene flow appear so weak? Here, we leverage this exceptional opportunity to address whether and how divergent floral syndromes in parent species shape gene flow across the hybrid zone. Using phenotypic and genomic data from two separate *P. davidsonii*–*P. newberryi* hybrid zones in the eastern Sierra Nevada, we answer the following questions: (1) How genetically divergent are the two parent species and what is the genetic architecture of species barriers? If gene flow occurs between parent species, we predict reduced overall divergence but for genomic differentiation to be exacerbated in narrow windows underlying key traits that diagnose species. If few loci underlie species identity, we expect genomic islands of divergence in an otherwise homogenized genome. Alternatively, a highly polygenic basis to species identity may generate continents of divergence. (2) Which genomic regions represent barrier loci, and to what degree are loci responsible for adaptation to pollinators barriers? If floral differentiation is important for maintaining species boundaries, we expect loci underlying species-diagnostic floral traits to be fixed or nearly fixed in parent species and to be identified as strong reproductive barriers.

## Results

### Continuous variation in hybrid ancestry is structured by elevation

We obtained whole-genome resequencing data and floral phenotypic measurements for 358 samples across two hybrid zones, Virginia Lakes (n = 115) and Gem Lakes (n = 243), in the eastern Sierra Nevada (Figure 1a–c). We mapped sequencing reads to the *P. davidsonii* reference genome based on its high quality annotation (Ostevik et al., 2024) and minimal observed reference genome bias (see Supplemental Materials for a detailed comparison of reference genome mapping rates). Prior to applying filtering thresholds, we identified 56,279,850 SNPs; after all filtering thresholds were applied, our genomic dataset contained 7,322,630 SNPs, with mean depth on the eight largest pseudochromosomes ranging from 4.44x – 5.37x, and mean coverage ranging from 71.26% – 79.27% (Tables S2–S3).

**Figure 1.**
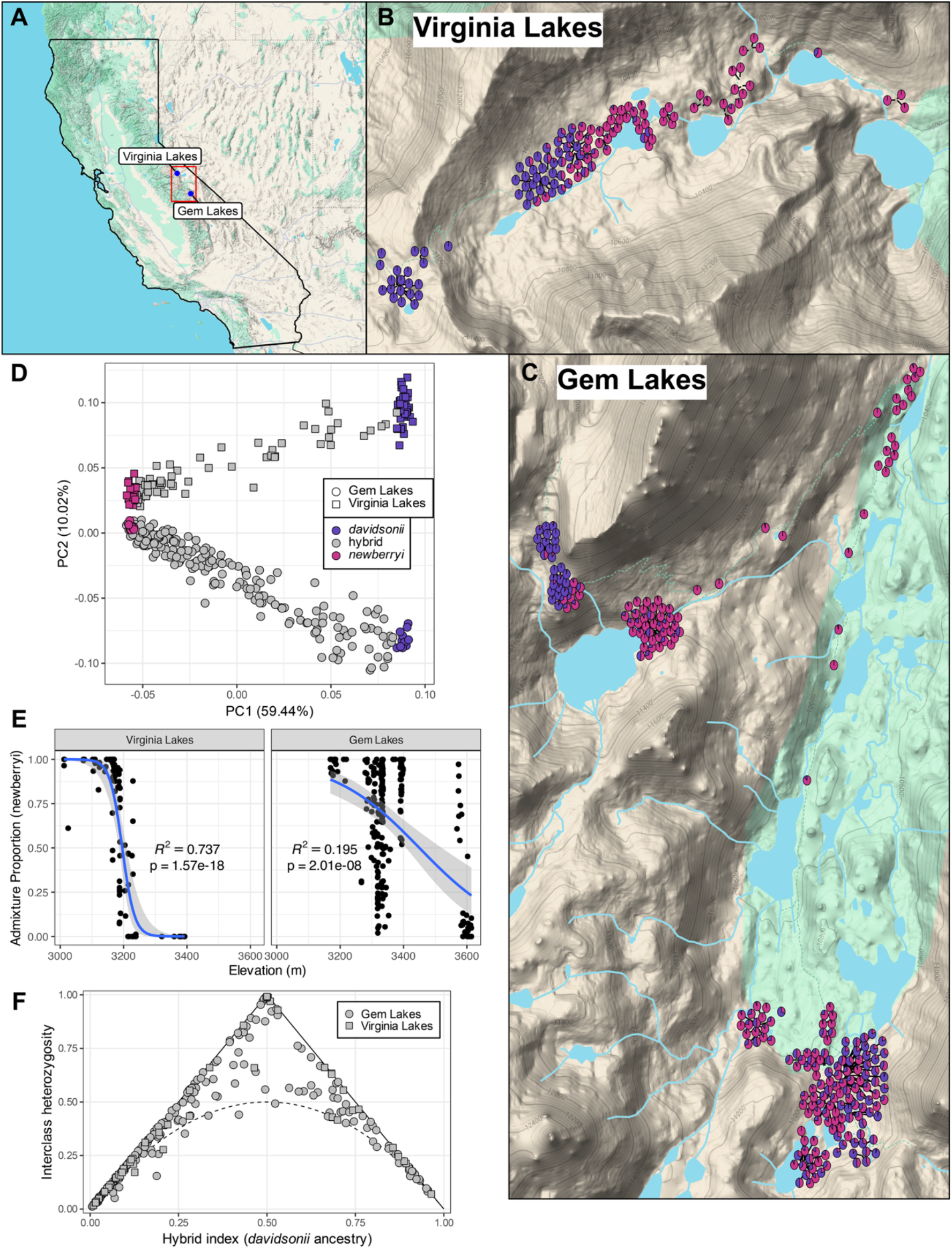
Overview of the hybrid zones. (A) Locations of the two focal hybrid zones within the Sierra Nevada. (B–C) Locations of all individuals used in this study; pie colors represent admixture proportion for *P. davidsonii* (purple) vs. *P. newberryi* (pink) at both sampling locations. (D) Genomic PCA constructed in PLINK2. (E) Logistic regression between admixture proportion and elevation. (F) Triangle plot of hybrid index and interclass heterozygosity for admixed individuals. Values are derived from local ancestry inference. The dashed curve marks the maximum heterozygosity expected under Hardy-Weinberg equilibrium for a single generation of admixture.

We found strong evidence of population structure, with the first axis of the genomic PCA distinguishing *P. davidsonii* from *P. newberryi*, and the second axis distinguishing the Virginia Lakes population from the Gem Lakes population (Figure 1d). PC2 tracks differentiation between *P. davidsonii* populations more strongly than between *P. newberryi* populations, suggesting there may be less gene flow between high-elevation *P. davidsonii* populations than between lower elevation *P. newberryi* populations. There was a broad range of intermediate PC1 and PC2 values, indicating a continuum of admixed ancestry in both hybrid zones. Our Admixture (Alexander et al., 2009) and local ancestry (Corbett-Detig & Nielsen, 2017) analyses, which were performed on LD-pruned subsets of 893,961 (no imputation) and 44,095 SNPs (after imputation), respectively (Table S3), revealed ongoing hybridization and potential introgression in both hybrid zones (Figure 1b–c, f). Admixture bar plots of K = 1–5 and the CV error curve are presented in Figures S2–S3. The minimum CV errors were found for K=2 and K=3, with K=3 slightly lower than K=2 (Δerror = 0.001). Given the known biology of the system (i.e., two parental species) and the negligible improvement from K = 2 to K = 3, we focus on K = 2 throughout the study. For our local ancestry analyses, while the majority of admixed individuals appear to be advanced generation backcrosses with either parent species, there are still a considerable number of early generation hybrids in both hybrid zones (Figure 1f). We find no obvious support for an earlier hypothesis from Kimball (2008) that hybrids primarily backcross with *P. newberryi*: the greater frequency of individuals with majority *P. newberryi* ancestry likely reflects our limited sampling at the highest elevations where majority *P. davidsonii* hybrid individuals would more likely be found. Major parent ancestry was strongly associated with elevation at both sites; individuals harbored increasingly larger proportions of *P. davidsonii* ancestry at higher elevations, consistent with parent species’ elevational distributions (Figure 1e). This pattern was particularly pronounced at Virginia Lakes, where sampling spanned an approximately linear elevational transect and hybrid individuals were largely restricted to a narrow band near 3200 meters (10,500 feet). The association with elevation was still present but less pronounced at Gem Lakes, where samples were collected across a more heterogenous landscape (Figure 1e).

### Parent species display genomic “continents”, rather than “islands” of divergence

Scans of the genomic landscape of divergence between parent species revealed heterogeneous patterns of genomic differentiation. Genome-wide dxy was 0.013 (5th – 95th percentiles: 0.0059 – 0.0181), genome-wide π for *P. davidsonii* was 0.0065 (5th – 95th percentiles: 0.0023 – 0.0129), and genome-wide π for *P. newberryi* was 0.0072 (5th – 95th percentiles: 0.0031 – 0.0128). Quantiles for each metric are presented in Table S4. In contrast to localized barriers to reproduction reflected in genomic islands of differentiation, we find that large tracts of the genome are elevated in both dxy and FST, consistent with a pattern of continents of divergence (sensu Michel et al., 2010) (Figure 2a; Figure S4 for no smoothing). These relatively highly differentiated regions coincide with increased genic content and decreased π in both parent species. Genome-wide, FST and dxy were positively correlated (Figure 2b), whereas FST was negatively correlated with π, and dxy was weakly positively correlated with π (Figure S5). In gene-poor genomic windows, π was similar between the two parent species, but it declined with increasing gene density in both species. This decline was more pronounced in *P. davidsonii* than in *P. newberryi*, resulting in lower π in *P. davidsonii* in gene-rich genomic windows (Figure 2c).

**Figure 2.**
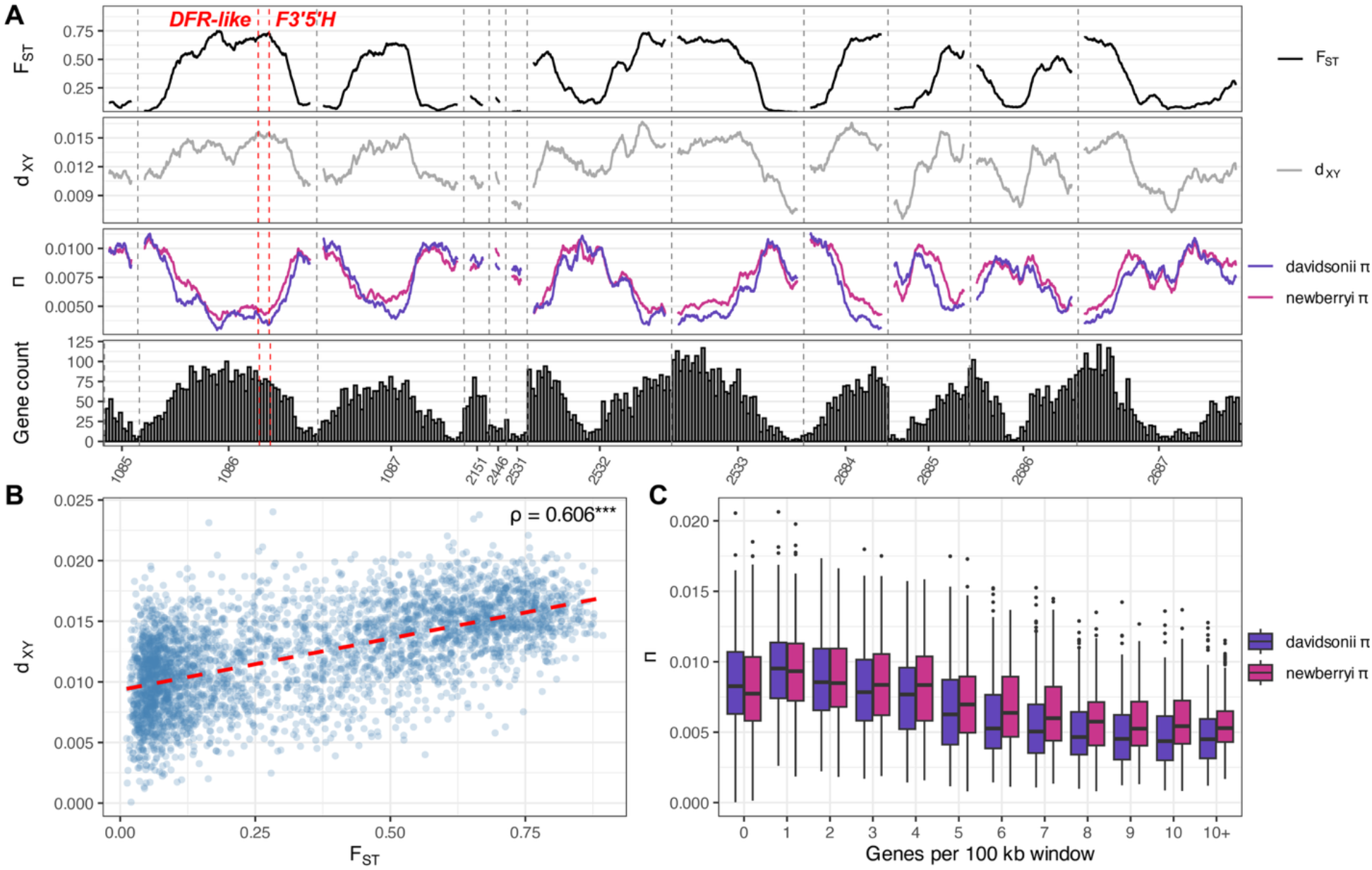
(A) Genome-wide patterns of FST, dxy, nucleotide diversity (π), and gene count in 100kb non-overlapping windows between “pure” *P. davidsonii* and *P. newberryi*. Values of dxy, FST, and π are smoothed across 40 100kb windows as a rolling mean. Gene count histograms are based on annotations for the *P. davidsonii* reference genome. (B) Genome-wide correlation analysis for FST vs. dxy between parent species in 100kb windows. Pearson’s ⍴ with *p* < 0.001 is denoted with three asterisks. (C) Boxplots of nucleotide diversity for each parent species, grouped by the number of genes per 100kb genomic window.

### Floral trait variation confirms divergent pollination syndromes in parent species

While the flowers of parent species are readily distinguishable by eye (Figure 3a), we quantified floral trait variation in both parent and hybrid samples across both hybrid zones using a Random Forest (RF) classifier. Traits included in the RF classifier include floral hue and chroma, external corolla length and width measurements, and internal length and width measurements including stamens, styles, and nectaries; the entire list of traits, and their biological relevance, is included in Table S5. The RF classifier perfectly assigned individuals to their parent species (OOB error = 0%; Table S6), confirming strong morphological differentiation. The traits most informative for classification, based on the Gini index, included floral hue and chroma (ranked 1st and 5th in the RF classifier), stamen exsertion (4th), and height of the floral opening (6th), all classic features of pollination syndromes (Figure S6). The RF classifier also highlighted additional traits not traditionally associated with pollination syndrome, including constriction and inflation near the base of the floral tube (2nd and 3rd). These traits might function to regulate access to nectar that is produced at the very proximal end of the floral tube. Other traits, such as floral tube length and mouth constriction, further distinguished parent species, but were less species-diagnostic (Figure 3c). A PCA of the most informative traits corroborated these results: floral hue and chroma, stamen exsertion, tube base constriction/inflation, and overall tube length loaded strongly on PC1, which explained >50% of total variance (Figure 3b). Mouth constriction and floral width traits loaded primarily on PC2, reflecting variation within, rather than between species. Although parent species’ flowers were clearly phenotypically differentiated, hybrid flowers were generally intergrade and formed a continuum for most traits (Figure 3b–c), including those not identified as the most strongly species-diagnostic (Figure S7).

**Figure 3.**
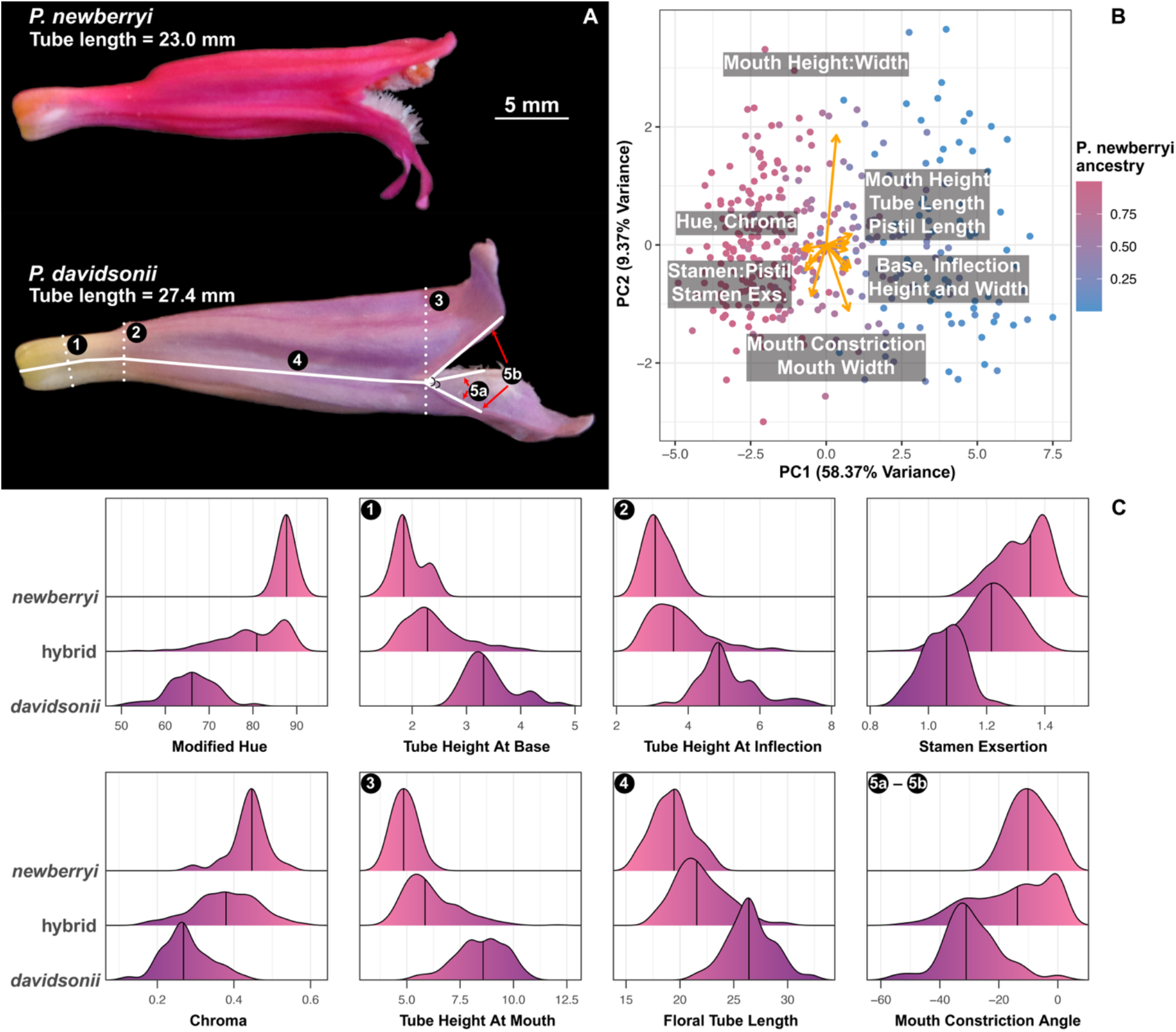
Quantification of floral differentiation between parent species. (A) Profiles of a typical *P. newberryi* and *P. davidsonii* flower, to scale. Lines and numbers on the *P. davidsonii* flower correspond to some of the key traits measured from the profile photographs. (B) PCA of floral trait multivariate space. Points are colored according to their proportion of *P. newberryi* ancestry, as inferred with Admixture. (C) Ridgeline plots of the top eight most informative traits as identified with the Random Forest (RF) classifier. Plots are ordered from top-to-bottom and left-to-right in order of importance in the RF classifier. Numbers on plots correspond to measurements in panel A.

### Floral trait variation is explained by genome-wide ancestry

We examined relationships among floral phenotypic variation (floral PC1), admixture proportion, and elevation using generalized linear models (GLMs). Floral phenotype was strongly predicted by genome-wide ancestry in both the Virginia Lakes (R2 = 0.868; p < 0.001) and Gem Lakes (R2 = 0.726; p < 0.001) sites (Figure 4); as flowers become more phenotypically similar to either parent species, their genomes also more closely resemble that species. We also detected significant, though weaker, relationships between floral phenotype and elevation: plants at lower elevations were more similar to *P. newberryi*, and plants at higher elevations were more similar to *P. davidsonii* (Figure 4), at both the Virginia Lakes (R2 = 0.375; p = 0.001) and Gem Lakes (R2 = 0.0886; p < 0.001) sites. These elevational trends echo Kimball (2008) and indicate that geographic variation in floral traits largely reflects shifts in hybrid ancestry across the contact zones. Consistent with Kimball (2008), although hybrids overall are phenotypically intergrade and occupy a continuum of trait-space between both parent species, more hybrid individuals skew toward a *P. newberryi*-like phenotype rather than a *P. davidsonii*-like phenotype.

**Figure 4.**
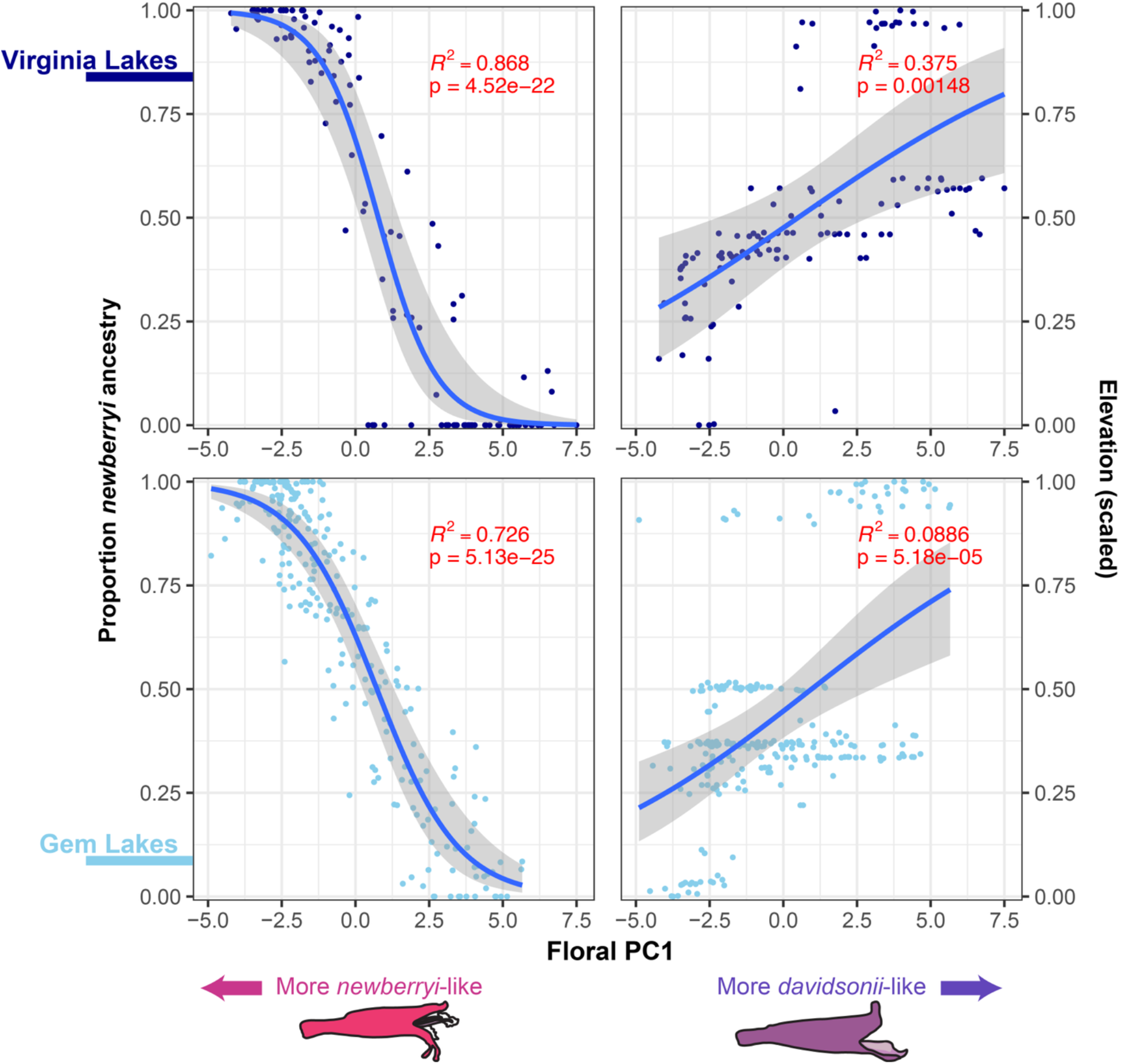
Results of generalized linear models (GLMs) testing the relationship between floral PC1 and admixture proportion (left) and elevation (right) at the Virginia Lakes (top) and Gem Lakes (bottom) hybrid zones. Elevation values are scaled from 0 to 1 within each hybrid zone.

### Broad association peaks reveal the genetic basis of key floral traits

We performed genome-wide association studies (GWASs) to identify loci underlying variation in species-diagnostic floral traits. After removing results with <5 SNPs of support per pseudochromosome, we identified significant associations in ten genomic regions across seven traits (Table 1). We found significant association peaks for two species-diagnostic traits: (1) floral hue and (2) tube height at inflection point. We also discuss results for a third trait, stamen length ratio. Results for additional traits are presented in Figures S8–S12.

**Table 1.** GWAS summary.

| Trait | Scaffold | Min bp | Max bp | SNP count | % Fixed SNPs | Cline |
| --- | --- | --- | --- | --- | --- | --- |
| Crease to Bottom Height | 1087 | 13,790,155 | 15,424,588 | 6 | 0.00% | Neutral |
|  | 2532 | 34,082,307 | 44,416,760 | 37 | 16.22% | Shallow |
|  | 2687 | 139,020 | 5,368,160 | 10 | 10.00% | Steep |
| Modified Hue | 1086 | 34,150,604 | 46,470,553 | 17575 | 64.48% | Shallow |
| Mouth Angle | 1086 | 37,196,033 | 42,282,447 | 98 | 3.06% | Shallow |
| Platform Angle | 2532 | 32,963,117 | 44,710,965 | 2330 | 60.09% | Shallow |
|  | 2686 | 7,993,759 | 8,135,661 | 23 | 0.00% | Neutral |
| Stamen Length Ratio | 2532 | 26,286,960 | 30,179,846 | 188 | 9.04% | Shallow |
| Tube Height at Inflection | 2532 | 37,854,627 | 42,260,154 | 7 | 14.29% | Shallow |
| Upper Petal Angle | 1086 | 35,143,450 | 43,349,327 | 222 | 73.87% | Shallow |

Significance assessed using permutation threshold. Only includes peaks with > 5 SNPs of support. % Fixed SNPs are the percentage of GWAS significant SNPs with AFD = 1 between parent populations. GWAS windows were assigned "shallow" or "steep" if the sliding window cline slope proportion (see genomic clines section) exceeded 10% for shallow or steep clines, respectively, at any point in the GWAS interval.

We identified a single major association peak for floral hue that encompasses a large genomic region on pseudochromosome 1086, spanning more than 12.3 Mb, and containing more than 17,000 SNPs exceeding the genome-wide significance threshold (Table 1; Figure 5). This region contains the anthocyanin pathway gene flavonoid 3’,5’ hydroxylase (*F3’5’H*), which catalyzes hydroxylation of the B-ring of flavonoids to produce blue delphinidin-type anthocyanins, and has been repeatedly implicated in floral color transitions in *Penstemon* and other angiosperms (Rausher, 2008; Stone & Wessinger, 2024; Wessinger & Rausher, 2015). The bluish-flowered *P. davidsonii* produces delphinidin-type anthocyanins in petal tissues, whereas the magenta-flowered *P. newberryi* produces cyanidin-type anthocyanins – a shift that can be accomplished by inactivation of the *F3’5’H* gene in *P. newberryi*. In fact, we previously found that the *F3’5’H* sequence in *P. newberryi* has a deletion of the second exon (Stone and Wessinger 2024). Here we found four SNPs surpassing the significance threshold located in the genic and putative promoter region of *F3’5’H*. There were several peaks in dxy 5-7kb upstream of *F3’5’H*, but no clear pattern of increased nucleotide diversity for *P. newberryi* or *P. davidsonii* at the *F3’5’H* region (Figure 5b).

**Figure 5.**
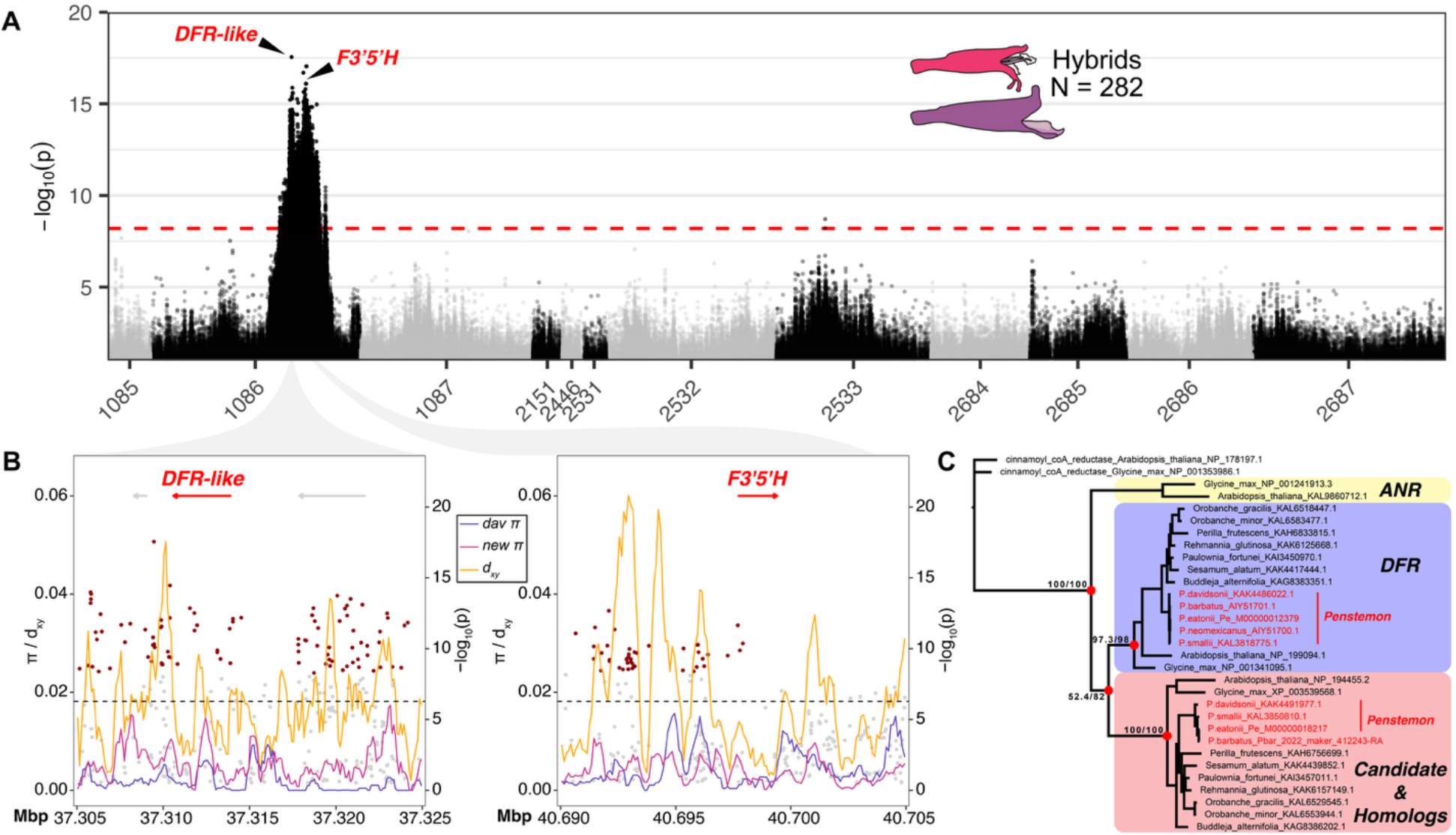
GWAS of floral hue. (A) A Manhattan plot for GWAS of floral hue variation identified one large association peak. Association support is plotted as −log10(*p*wald). The dashed red line denotes the genome-wide significance threshold as determined by permutation. (B) Zoomed-in plots of two focal candidate genes. Focal genes within each window are annotated. The dashed black line denotes the genome-wide 95th percentile of dxy. SNPs exceeding the genome-wide GWAS significance threshold are in red. Values of π and dxy were estimated in 100bp windows, smoothed in 5-window rolling means prior to visualization. (C) A gene tree including the *DFR-like* candidate gene. Clades for *DFR*, *ANR*, and the *DFR-like* candidate gene (including putative homologs) are annotated. The gene tree is rooted with a Cinnamoyl-CoA reductase gene in *Arabidopsis thaliana.* Red dots at nodes highlight focal relationships of interest, and text at nodes represents SH-aLRT support (%) / ultrafast bootstrap support (%). Tips in red indicated *Penstemon* samples.

While the presence of *F3’5’H* in our major floral hue GWAS peak strongly suggests its involvement, the SNP most strongly associated with floral hue (chr1086:37309460; *p* = 2.79e-18) was located approximately 3.39 Mbp away from of *F3’5’H*, and 1131 bp downstream of a gene annotated as dihydroflavonol 4-reductase (*DFR*) (Figure 5a). In plants, *DFR* is a key enzyme in the anthocyanin biosynthetic pathway, catalyzing the NADPH-dependent reduction of dihydroflavonols to leucoanthocyanidins. This step directs metabolic flux toward the production of anthocyanins and proanthocyanidins, which are major determinants of floral pigmentation and can influence traits such as hue, intensity, and patterning (Saito et al., 2013). We constructed a gene tree including this gene model, a second *P. davidsonii* gene model with high similarity to annotated *DFR*s, and representative *DFR* and anthocyanidin reductase (*ANR*) protein sequences from a broad sampling of angiosperms. This analysis revealed that the *DFR* gene found within our association peak belongs to a clade that is paralogous to most well-studied *DFR* genes, making it difficult to speculate on function (Figure 5c). Within this "*DFR-like*" gene sequence and in its putative promoter region (<1kb upstream), we identified an additional 13 SNPs exceeding the genome-wide significance threshold. This region was also characterized by relatively high dxy and a relative increase in nucleotide diversity in *P. newberryi* (Figure 5b).

The association peak for floral tube height at the inflection point spanned more than 4.4 Mbp on pseudochromosome 2532 and contained 7 SNPs exceeding the genome-wide significance threshold (Table 1; Figure S13a). The most strongly associated SNP (chr2532:40638154; *p* = 9.54e-10) was located within the genic region of *CYP707A2*. One additional SNP exceeding the genome-wide significance threshold fell within the *CYP707A2* gene model. Members of the *CYP707A* family encode abscisic acid (ABA) 8’-hydroxylases, cytochrome P450 enzymes which catalyze the primary step in ABA catabolism. ABA is a versatile plant hormone that regulates a wide range of processes, including seed dormancy and germination, stress responses, and vegetative growth. It also influences reproductive development, with roles in floral organ growth, flower opening, and senescence, highlighting its broad regulatory functions across both vegetative and reproductive tissues (K. Chen et al., 2020).

The association peak for stamen length ratio spanned nearly 4 Mbp on pseudochromosome 2532 and contained 188 SNPs exceeding the genome-wide significance threshold (Table 1; Figure S13b). The most strongly associated SNP (chr2532:28618399; *p* = 3.36e-17) was located in a repeat-dense region, with the closest annotated gene (a putative β-glucuronosyltransferase) > 12kb away. Other notable genes of interest in this association peak include three xyloglucan endotransglucosylase proteins (*XTH*; 2 SNPs within CDS), which regulate cell wall loosening and expansion (Ishida & Yokoyama, 2022; Rose et al., 2002), and two putative auxin efflux carrier component proteins (*PIN1C*), which control the direction and intensity of auxin flow and have been implicated in numerous aspects of plant development, including floral development (Cheng & Zhao, 2007; Cucinotta et al., 2021).

### Variation at candidate anthocyanin genes underlies floral hue and pollinator perceptibility

Floral hue was strongly related to petal pigment composition, with the proportion of delphinidin relative to cyanidin in floral tissues explaining 80% of the variation in hue (adjusted R2 = 0.80; p < 0.001) (Figure 6a). To evaluate the ecological consequences of this variation in floral hue and its potential contribution to reproductive isolation, we modeled color perception for the two primary pollinators in the *P. davidsonii*–*P. newberryi* hybrid zones: hummingbirds and bees. Under these visual models higher just-noticeable difference (JND) values indicate greater detectability. For the hummingbird visual model, mean JND values were 7.4, 9.8, and 11.3, for *P. davidsonii*, hybrids, and *P. newberryi*, respectively, with all pairwise differences significant in post hoc Tukey tests (Figure 6b). In contrast, mean JND values for honeybees (7.4, 7.6, and 7.4, respectively) did not differ significantly among groups (Figure 6b). Thus, differences in floral hue, which itself is a product of pigment composition, correspond to differences in conspicuousness for hummingbirds, but not for bees. Consistent with this, flowers occupy overlapping regions of honeybee color space (Figure 6c), suggesting that although bees can distinguish flowers from a green background, they may be unable to differentiate parent and hybrid flowers by color alone. This pattern supports earlier work by Kimball (2008), which implicated small bees as the most likely vectors of gene exchange in this system.

**Figure 6.**
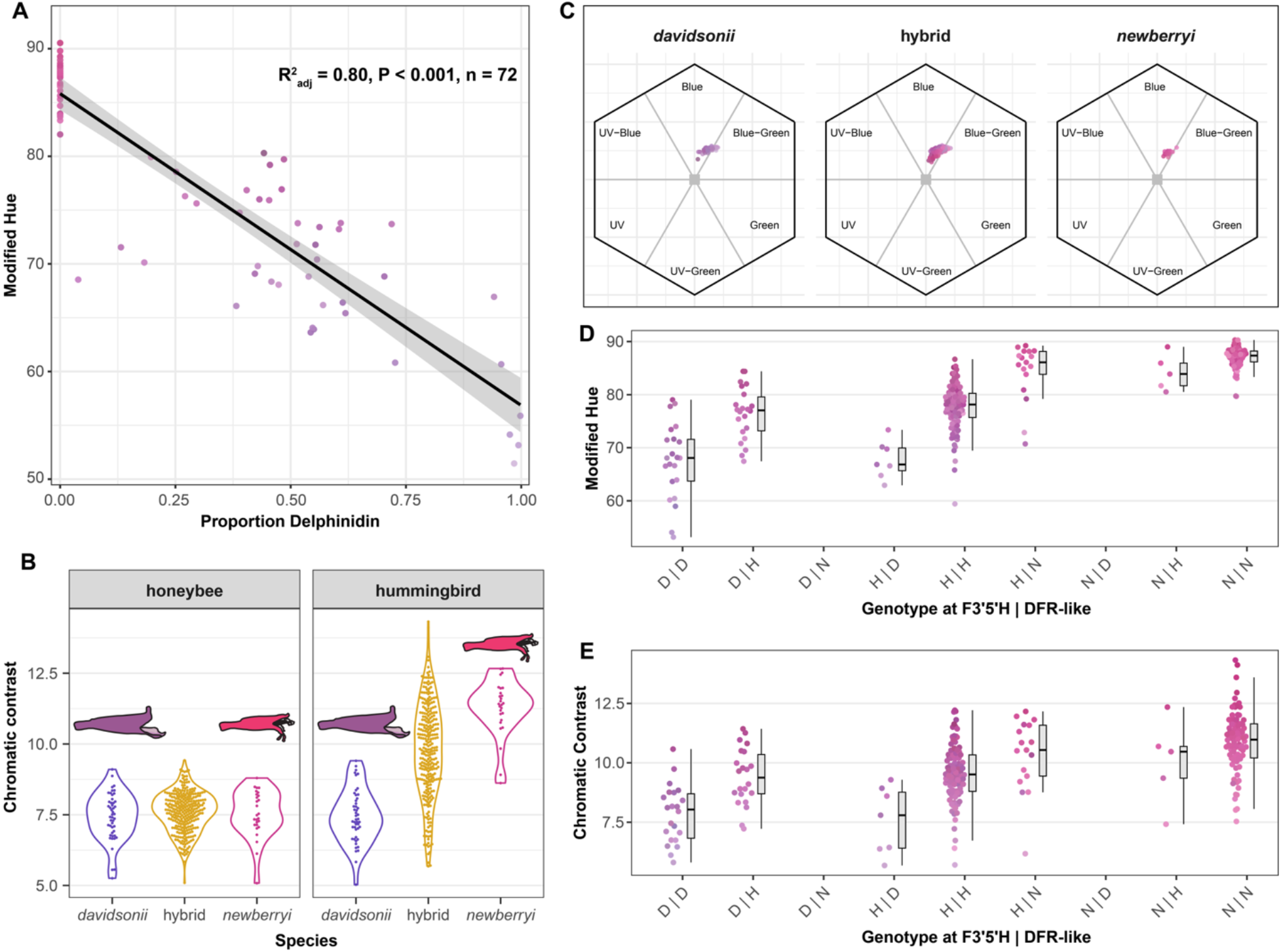
The molecular basis of pigmentation and its effect on pollinator discernibility and floral hue. (A) Linear regression between the proportion of delphinidin in floral tissues and floral hue. (B) Chromatic contrast (JNDs) of honeybee vs. hummingbird viewers of parental and hybrid flowers. When JND = 1, hues are perceptible against a leaf background, with JND > 1 implying increasingly distinct hues. (C) Bee color vision models in hexagonal color space for *Penstemon davidsonii*, hybrids, and P. *newberryi*. Each point represents a flower’s position based on reflectance spectra transformed into bee visual space, with axes corresponding to UV, blue, and green photoreceptor stimulation. Values of (D) floral hue and (E) hummingbird chromatic contrast with respect to genotype at *F3’5’H* and the *DFR-like* candidate genes within the hue GWAS association peak.

To further investigate the genetic architecture for floral hue and pigmentation, we characterized patterns of variation in genotype on floral hue and pigment composition at the two anthocyanin candidate loci within the floral hue GWAS association peak: *F3’5’H* and the *DFR-like* candidate gene. Because these loci were identified within the GWAS peak using the same individuals, the results described below characterize potential interactions between candidate loci within the GWAS peak rather than representing an independent test of association with traits of interest. Three nested linear models compared floral hue with ancestry at these loci. The additive model, in which *F3’5’H* and *DFR-like* each contribute independently, was best supported, suggesting that both genes may influence hue but without evidence for epistasis (Table S7). Mean hue values across ancestry classes indicate that *DFR-like* is most strongly associated with hue when *F3’5’H* is heterozygous (Figure 6d). Patterns of pigment production parallel those of hue: delphinidin was absent in individuals with *P. newberryi* ancestry at both loci and consistently present when both loci carried *P. davidsonii* ancestry (Figure S14). Likewise, patterns of hummingbird chromatic contrast vs. genotype mirror results for floral hue: individuals homozygous for *P. davidsonii* ancestry have lower chromatic contrasts, homozygotes for *P. newberryi* ancestry have the highest chromatic contrasts, and heterozygotes are intermediate (Figure 6e). Our interpretation is complicated by the lack of recombination between our candidate loci – no individuals carried opposite homozygous parental ancestry at the two genes – raising the possibility of linked selection. Thus, while ancestry at *F3’5’H* covaries with floral hue and pigment composition, it alone does not fully explain variation in these traits, and the contribution of the *DFR-like* locus remains unresolved.

### Loci underlying floral trait divergence are not barriers to gene flow

We fit clines in the probability of locus-specific ancestry along a genome-wide admixture gradient (i.e., genomic clines), as implemented in *bgchm* (Gompert et al., 2024). The main goals of the genomic cline analyses were to find genomic regions particularly resistant (i.e., “steep” clines) or permissive (i.e., “shallow” clines) of gene flow, as these regions are expected to inform on the genetic basis of reproductive isolation. Under the hypothesis that pollination syndrome traits contribute to reproductive isolation, we predict regions associated with syndrome traits should have steep clines. Cline slope and center standard deviations were estimated to be 0.129 and 0.632, respectively (Table S8). Of the 49,019 clines estimated, 6,917 (14.1%) were credibly steep (cline slope 95% confidence intervals > 1), indicating potential barrier loci, and 4,251 (8.67%) were credibly shallow (cline slope 95% confidence intervals < 1), indicating a potential lack of barriers. Three pseudochromosomes (2533, 2684, and 2687) were significantly enriched for steep clines (1,000 permutations: *p* < 0.001; Table S9), while an additional three (1086, 2151, and 2532) were significantly enriched for shallow clines (1,000 permutations: *p* < 0.001; Table S9). Notably, pseudochromosomes 1086 and 2532, which are enriched for shallow clines, contain nearly all significant GWAS association peaks (Table 1).

While we identified many steep clines across the genome, the region encompassing the major floral hue GWAS association peak has the largest excess of shallow clines in the genome (Figure 7a), representing an outlier compared to the genomic background (p < 0.001 with a Mann-Whitney U test; Figure S15a). Additionally, in contrast to the genomic background, which has cline centers distributed symmetrically around 0.5, cline centers in the hue association peak are significantly shifted toward *P. davidsonii* ancestry (Figure S15b). These results point to relaxed barriers to gene flow at the floral hue locus. Thus, while we identified many steep clines representing potential barrier loci across the genome, these steep clines are not associated with loci underlying floral syndrome traits.

**Figure 7.**
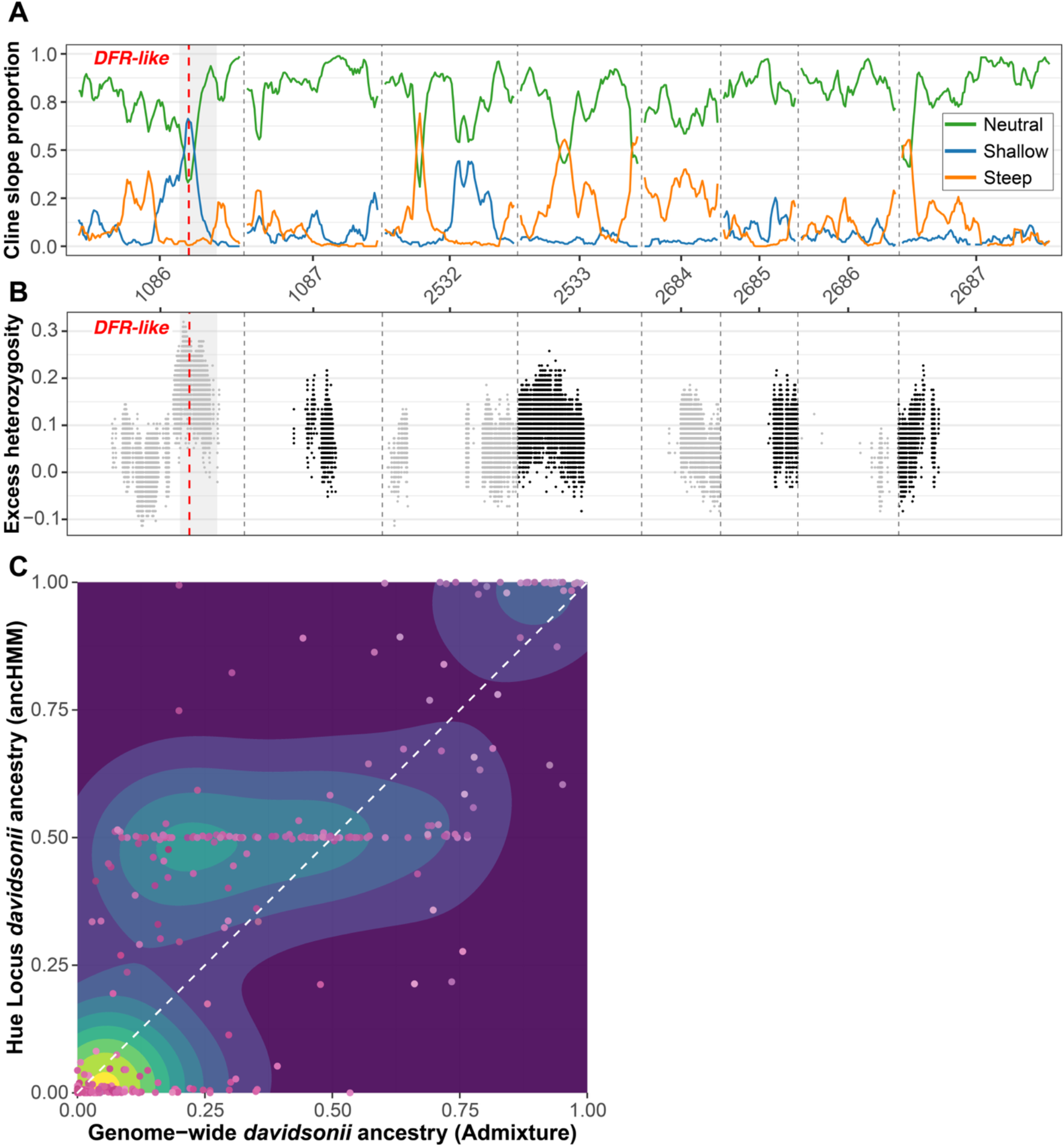
Genomic clines and patterns of heterozygosity. (A) Sliding window means of the proportion of clines with neutral, shallow, and steep slopes. Clines are credibly steep or shallow if their 95% probability distribution exceeds or is less than 1, respectively; otherwise, they are neutral. Cline proportions are averaged in 500kb windows and smoothed in increments of 5 windows. (B) Patterns of excess heterozygosity across the genome on hybrids with intermediate admixture proportions (25-75%), calculated at SNPs that are fixed between parent species (AFD = 1). (C) 2D kernel density estimates showing the relationship between genome-wide ancestry and ancestry at the floral hue GWAS association peak. Warmer colors on the density plot represent regions with higher estimated probability density, indicating a greater concentration of points in those areas. Each point is an individual, and points are colored by their floral hue. The white dashed line corresponds to the expected relationship between genome-wide ancestry and ancestry at the hue association peak.

We examined patterns of heterozygosity and local ancestry to better understand the nature of the shallow clines at the floral hue association peak. We calculated observed and expected heterozygosity, and excess heterozygosity in hybrids for each fixed SNP between parent species. The floral hue association peak had estimates of excess heterozygosity higher than any other genomic region, with observed heterozygosity in hybrids greater than 80% and a global peak in heterozygosity nearly 2Mbp downstream of our *DFR-like* candidate gene (Figure 7b). Using local ancestry inference, we found hybrids with a broad range of genome-wide ancestry (from majority *P. davidsonii* to majority *P. newberryi*) are completely or nearly completely heterozygous at ancestry-informative markers in the floral hue association peak (Figure 7c). This indicates that the shift in cline centers at this region are driven by excess heterozygosity, rather than excess *P. davidsonii* ancestry, *per se*.

## Discussion

### Despite decades of hybridization, species boundaries appear largely intact

Divergent ecological selection on traits that influence fitness and mating can strengthen reproductive isolation and promote ecological speciation (Nosil, 2012). Hybrid zones resulting from interbreeding between divergent taxa provide a window into these mechanisms and can reveal targets of selection in the genome that underlie ecological divergence. These ecological sources of selection are diverse and can include, for example, sexual selection (Dougherty & Carling, 2024; Long et al., 2024; Schield et al., 2024), pollinator-mediated selection (Campbell et al., 2024), precipitation/climate (Farnitano et al., 2025; Semenov et al., 2025; Sianta et al., 2024; Tataru et al., 2025), or migratory cues (Blain et al., 2025; Justen et al., 2024, 2025).

The genetic architecture of underlying traits shapes patterns of genomic divergence in hybridizing species. For example, relatively few barrier loci might present as obvious and narrow islands of elevated divergence relative to a homogenized background, while a highly polygenic basis to reproductive isolation might present as a genome-wide pattern of elevated divergence (i.e., continents). Yet heterogeneous genomic divergence can also arise in the absence of gene flow, where peaks of divergence reflect variation in diversity and linked selection (selective sweeps, background selection) rather than differential introgression (Cruickshank & Hahn, 2014; Harrison & Larson, 2016; Ravinet et al., 2017; Shang et al., 2023).

Our results show that genomic divergence and differentiation in the *P. davidsonii*–*P. newberryi* hybrid zone is elevated across broad, gene-dense regions of the genome, rather than in a few narrow islands associated with conspicuous floral traits. Both absolute divergence and relative differentiation are substantial and widespread, with multiple pseudochromosomes exhibiting extended regions of elevated dxy and FST (Figure 2a). The positive genome-wide correlation between FST and dxy (Figure 2b) indicates that highly differentiated genomic regions also tend to exhibit high absolute divergence, suggesting these regions are truly distinct rather than simply reflecting local reductions in nucleotide diversity (Cruickshank & Hahn, 2014). That the most highly divergent genomic regions are also the most gene-dense suggests an important role of linked selection and a general trend of selection against introgression in this system, as observed in other systems (e.g., Shang et al., (2023)). Genome-wide, we observed a decline in nucleotide diversity with increasing gene density, especially in *P. davidsonii* (Figure 2c). This pattern is consistent with the expectation that linked selection is strongest in gene-rich regions of the genome and may suggest that selection against introgression is stronger in *P. davidsonii* than in *P. newberryi*.

The co-occurrence of broad regions of elevated divergence with steep genomic clines (floral hue locus notwithstanding) suggests that many genomic regions resist gene flow and may act as effective reproductive barriers. That these regions are broad continents, rather than narrow islands, is more consistent with many differentiated (polygenic) barrier loci (Martin et al., 2019; Michel et al., 2010; Schumer et al., 2018; Todesco et al., 2020). Overall, our results are counter to the hypothesis that long-term hybridization between parent species has homogenized large portions of the genome, leaving only small islands of differentiation. To the contrary, we find that parent species have maintained large genomic regions of differentiation despite, despite a minimum of decades (Clausen et al., 1940) of secondary contact.

The mechanisms that produce continent-like patterns of divergence are varied. Inversions or other structural variants can suppress recombination across large genomic regions and thereby preserve linkage between many barrier loci (Stevison et al., 2011). Extended regions of low recombination magnify the effects of linked selection and allow divergence to accumulate. Polygenic selection acting on many small-effect loci may also produce broad differentiation without requiring a single large-effect gene. While we did not find evidence that structural variants produced these patterns, we also cannot rule out this possibility. Future investigations into this system should aim to explore the occurrence and distribution of structural variants with respect to putative barrier loci and loci underlying variation in key traits.

### Floral divergence is unlikely to appreciably confer reproductive isolation

Flowers are recurrent targets of divergent ecological selection because of the dual role of pollinators as vectors of gene flow and agents of selection on floral morphologies. In hybrid zones, lower visitation rates and/or efficiency of pollen transfer in plants with intermediate floral traits may reduce gene flow via pollinator-mediated divergent selection (sensu Waser & Campbell, 2004). This pollinator-mediated divergent selection is thought to be a major driver of floral phenotypic change and indeed ecological speciation in angiosperms (Kay & Anderson, 2025; Van Der Niet et al., 2014), including between bee- vs. bird-pollinated flowers (Campbell et al., 2024; Wessinger, 2024).

Our study system appears to be a classic case of floral isolation, with obvious differences in floral morphologies consistent with adaptation to divergent pollinators. The most important floral difference – petal hue – has a simple genetic architecture and maps to a single genomic region. If pollinator preference imposed strong reproductive isolation between parent species, we would expect to find strongly divergent genomic regions corresponding to variation in floral traits that resist introgression (i.e., show steep genomic clines). We predicted that the floral hue region in particular would present as a barrier locus exhibiting steep genomic clines if floral isolation is a primary reproductive isolating barrier. In fact, we found an abundance of shallow genomic clines and excess heterozygosity at the floral hue association peak (Figure 7a–b), suggesting this region is permissive of introgression and may even be under selection to maintain heterozygous ancestry in hybrids. Beyond the floral hue locus, we found that other loci generating genetic effects on pollination syndrome traits most often displayed shallow genomic clines (Table 1).

While many regions of the genome are resisting the homogenizing effects of gene flow and exhibit steep genomic clines, these barrier loci do not overlap with the genes conferring divergent floral adaptation. As a result, we conclude that floral divergence is unlikely to be a primary reproductive barrier in the *P. davidsonii*–*P. newberryi* hybrid zone, and suggest that other ecological or genetic factors must underlie the maintenance of species boundaries. Similar patterns have been documented in at least one other hybrid zone between plant species with clear floral differentiation (Stankowski et al., 2023), and indeed, this idea was proposed in the *P. davidsonii*–*P. newberryi* system already (Kimball et al., 2008; Kimball & Campbell, 2009), suggesting that reproductive isolation in these systems can be maintained by polygenic and/or postzygotic selection unrelated to floral divergence. However, the patterns described here contrast with data from a different species pair in *Penstemon* (the *P. barbatus–P. virgatus* species complex), where genomic differences between bee and hummingbird syndrome species map to narrow regions that correspond to floral trait QTLs (Wessinger et al., 2023). This suggests that the genomic architecture of species divergence can differ drastically, even among close relatives.

Although the two parent species show striking multi-trait floral differences, these trait differences may not generate strong enough assortative mating to prohibit gene flow. Secondary pollinators can contribute substantially to reproductive success by buffering plants against the absence or unreliability of primary pollinators (e.g., Wenzell et al., 2024). Indeed, even apparent specialist flowers are increasingly viewed as maximizing fitness through the use of suites of pollinators rather than strict dependence on a single most-effective pollinator (Kay & Anderson, 2025). Previous work in the *P. davidsonii*–*P. newberryi* hybrid zone identified considerable overlap in floral visitors between parent species and hybrids (Kimball, 2008). This result was corroborated by our pollinator visual models showing that bees may not be able to distinguish among parental and hybrid floral colors using chromatic contrast (Figure 6). Together, these observations indicate that secondary pollinators likely play a larger role in this system than previously appreciated and that apparent pollination “specialization” masks considerable functional overlap.

Reproductive isolation in this system may instead arise from elevational adaptation. We found that genome-wide admixture proportion is tied to elevation, such that individuals at higher elevations increasingly have on average higher proportions of *P. davidsonii* ancestry (Figure 4), consistent with a tension zone dynamic where selection acts against genotypes displaced from their local ecological optima. The combination of a narrow, apparently stable hybrid zone, broad genomic continents, and a lack of clear barrier loci linked to genomic regions underpinning floral variation, implies that selection maintaining species boundaries is more likely to be ecological and polygenic (i.e., selection on multiple physiological, phenological, and life-history traits covarying with elevation) than related to floral divergence. In particular, water use efficiency and maximum photosynthetic rate, having already been implicated as potential targets of divergent selection in this system (Kimball & Campbell, 2009), represent key traits to investigate in future studies.

### Major floral hue locus exhibits elevated heterozygosity

Our GWAS recovered a broad, highly significant association for floral hue centered on a large region containing *F3’5’H*. *F3’5’H* is a well-established determinant of pigmentation across angiosperms (e.g., Hopkins & Rausher, 2011; Ishiguro et al., 2012; Smith & Rausher, 2011), including *Penstemon*, where loss-of-function mutations to this gene are predictably associated with redder pigmentation and transitions to hummingbird pollination (Stone & Wessinger, 2024; Wessinger & Rausher, 2015). A large deletion to the second exon of *F3’5’H* in the *P. newberryi* sequence (Stone & Wessinger, 2024) undoubtedly abolishes protein function. Although our association peak encompassed *F3’5’H*, the most strongly associated SNP lies several Mbp upstream near a putative short-chain dehydrogenase/reductase, which we refer to as *DFR-like* (Figure 5). Short-chain dehydrogenase/reductases form one of the largest NAD(P)(H)-dependent oxidoreductase families in plants and include several enzymes with well-documented roles in flavonoid and anthocyanin biosynthesis (Moummou et al., 2012). Changes to *DFR* function have previously been shown to underlie evolutionary shifts from delphinidin to cyanidin production through sequence changes causing substrate specificity (e.g., Smith et al., 2013), making this a plausible target for a shift in floral hue. Although hundreds of genes lie within the floral hue association peak, the presence of this second, mechanistically plausible gene suggests floral hue may be controlled by more than *F3’5’H*.

Inference of causality from our GWAS is limited by the broader genomic context. Given the size of the association peak and the sheer number of possible candidate genes included in the interval, strongly significant SNP associations could be driven by linkage to the true causal gene alone. The strong allele frequency differences observed between our parent species creates tracts of parental ancestry that co-segregate in hybrids (admixture LD), even when sampled across multiple generations. These features prevent fine-scale resolution of causal variants within the floral hue association peak. Ultimately, distinguishing among plausible genes in the floral hue association peak will require functional data such as expression analyses, enzyme assays, or transgenic validation. Therefore, while our results provide a clear framework for future work, demonstrating that a major-effect locus exists and identifying a novel candidate gene, they do not yet allow definitive identification of the causal determinants of floral hue in this system.

While our genomic cline analyses uncovered many steep clines scattered across the genome, the major floral hue association peak – our primary candidate for a barrier trait – showed a remarkable abundance of shallow clines and extreme heterozygosity (Figure 7). This elevated heterozygosity could in theory indicate selection for the intermediate floral hue carried by heterozygous individuals (Figure 6), imposed either by pollinators or by other agents of selection on flower color (Narbona et al., 2021; Strauss & Whittall, 2006). Linked selection on complementary partially deleterious haplotypes can also generate a pattern of elevated heterozygosity (pseudo-overdominance; Gilbert et al., 2020), raising the possibility that elevated heterozygosity has nothing to do with flower color. Although we currently cannot identify the source of excess heterozygosity, overall our results are inconsistent with a simple model in which a small number of barrier loci underpinning variation in floral traits confer reproductive isolation while the remainder of the genome freely homogenizes.

## Conclusions

We employed a combination of phenotypic and genomic analyses across two replicate *P. davidsonii*–*P. newberryi* hybrid zones in the eastern Sierra Nevada to understand the genetic architecture of trait variation and reproductive isolation. Parent species were strongly differentiated in floral morphology, including pigmentation, and we recovered clear suites of traits that reliably distinguish parent species. Despite at least decades of hybridization, genome-wide differentiation between parent species was substantial and was especially pronounced in gene-dense genomic regions. GWAS on species diagnostic floral traits identified a large hue-associated genomic region containing one previously identified (*F3’5’H*) and one novel (a short-chain dehydrogenase/reductase; *DFR-like*) candidate gene for pigmentation. Genomic cline analyses revealed many steep clines distributed throughout the genome, suggesting a polygenic basis for reproductive isolation. Yet, contrary to the expectation that floral trait loci generate barriers to gene flow, the floral hue association peak contained an abundance of shallow genomic clines, implying this region is quite permissive of introgression. Pollinator visual modeling indicates that while hummingbirds can distinguish among parent species floral hues, bees cannot, suggesting a potential mechanism for the continued production of hybrids and supporting the lack of steep clines at the floral hue locus. Excess heterozygosity at the floral hue association peak further supports this finding and suggests selection may be acting to maintain variation at this locus. Collectively, our results demonstrate a decoupling between conspicuous floral divergence and genomic barriers to introgression and suggest that despite apparent adaptation to different suites of pollinators in each species, floral differentiation does not act as a strong reproductive barrier; instead, many barrier loci are related to some unmeasured source(s) of selection.

## Materials and Methods

### Sampling

We collected leaf tissues and whole flowers from a total of 358 individual plants across two main contact zones (Virginia Lakes, n = 115; Gem Lakes, n = 243) in the eastern Sierra Nevada in the summers of 2022 and 2023. (Figure 1a–c). At each site, we sampled roughly along an elevational gradient, extending from lower elevations in the valley, where all plants phenotypically appear to be “pure” *P. newberryi*, through the contact zone at intermediate elevations, to high elevation sites above the tree line where all plants phenotypically appear to be “pure” *P. davidsonii*. Fresh leaf tissues were collected from plants and immediately placed into individual bags containing silica dessicant. Flowers were collected by clipping at the pedicel, placing into a 15 mL falcon tube, and storing on ice/cooling packs. We ensured collection of 1-2 flowers per plant, and only sampled flowers that had opened completely. Due to variation in flowering time between species and across samples in the hybrid zone, we collected flowers that were both pre-and post-anther dehiscence, but ensured all floral components were intact. Post-collection, flowers were wrapped in moist tissue paper and stored at 4°C for up to one day prior to phenotyping.

### Phenotyping

Whole flowers were pinned on a black background and photographed with a Canon EOS M50 Mark II with a scale in-frame. Flower petals were then removed to be processed for color (see below), and whole stamens and pistils were imaged using a Dino-Lite Premier digital microscope (Dunwell Tech.) set to 20x magnification with a scale in-frame. Stamens were further dissected and oriented to image nectaries, which are present at the base of the stamen and indicated by a discoloration of tissue; nectary area is correlated with nectar volume in other *Penstemon* species (Katzer et al., 2019). Floral traits other than color were then measured from images using imageJ (Abràmoff et al., 2004). When multiple measurements were available for a single sample (e.g., four nectaries across two flowers), they were averaged for a composite measurement. Trait measurements are reported in millimeters or mm2, to the nearest 1/10th millimeter. A full list of traits measured are listed in Table S5.

Flower color was measured via reflectance spectroscopy on the upper lobe of petals using an Ocean insight FLAME-S-UV_IS spectrometer equipped with a Y-shaped reflection probe and a deuterium/tungsten light source (DH-2000-BAL, Ocean Insight). Reflectance spectra were recorded from 300-800 nm in a dark room after calibration using a Spectralon diffuse reflectance standard (WS-1, Ocean Insight). Flower petals were sandwiched between two glass microscope slides and measured at a 45-degree angle relative to the reflectance probe to limit interference from floral curvature on reflectance measurements. Flowers were stored in silica gel following reflectance measurements. Raw floral reflectance spectra were imported into R and analyzed with *pavo* v.2.9.0 (Maia et al., 2019). Negative values were converted to 0, and reflectance spectra were averaged across flowers for samples with multiple flowers. These spectra were used to calculate color attributes (hue, chroma, and brightness) for each sample. We used a segment classification approach to calculate hue and chroma, and brightness was calculated as the total area under the curve from 400-700 nm, following Smith (2014). Hue in degrees was then calculated from 0° (magenta/red) to 360° (blue/purple), following Smith (2014). Because our calculated hue values bridged the 0° / 360° boundary, we report a “modified hue” value obtained by subtracting 270° from hues > 270° and adding 90° to values < 270°. This modification ensures flowers with redder hues always have higher values than flowers with bluer hues (as blue flowers all have hues > 270°) and avoids issues where red flowers may have hues near 360° or 0°.

### DNA extraction, sequencing, and QC

We extracted DNA from silica-dried leaf tissues using a modified CTAB protocol (Doyle & Doyle, 1987). We submitted DNA samples to the Duke University Sequencing and Genomic Technologies core facility for whole-genome Illumina library preparations using the Illumina Tagment DNA kit (Illumina Inc.) and sequencing to ∼5x depth using 150-bp paired-end reads generated on the NovaSeq X Platform (Illumina Inc.). Raw Illumina reads were quality filtered in the same manner as Stone and Wessinger (2024): filtering began with fastp v.0.23.2 (S. Chen et al., 2018), enabling auto-detection of adapters, limiting read length to 30 bp, removing unpaired reads, enabling base correction for overlapping reads, and enabling poly-x trimming on 3*’* ends of reads. We then verified final sequence quality with fastQC v.0.11.9 (Andrews, 2010) and multiQC v.1.10.1 (Ewels et al., 2016). Quality-filtered reads were then mapped to the annotated *P. davidsonii* reference genome (Ostevik et al., 2024) with bwa-mem v.0.7.17 (Li, 2013), and only reads passing a mapping quality threshold of *q* > 30 were retained. Duplicated reads were marked and removed with SAMtools markdup v.1.15.1 (Li et al., 2009), and overlapping paired-end reads were clipped with bamutil clipOverlap v.1.0.15 (Jun et al., 2015).

To determine whether the choice of reference genome may impact the results of our genomic analyses, we compared genome-wide mean read depth and coverage, and per-sample missing data rates (from a filtered .vcf), for all individuals when reads were mapped to the *P. davidsonii* (PRJNA1010203) (Ostevik et al., 2024) and *P. newberryi* (PRJNA985252) reference genomes. Both datasets were processed using identical pipelines for read filtering, mapping, variant calling, and variant filtering, as described above.

### Variant calling and filtering

Read assembly and variant calling was performed with bcftools v1.15.1 (Li, 2011). We used the multivariant caller and only called variants with BQ ≥ 20 and MQ ≥ 30. Indels were re-aligned with bcftools norm. After re-aligning indels, we calculated summary statistics for an initial “unfiltered” data set, and then applied additional filters with vcftools v0.1.17 (Danecek et al., 2011) and bcftools. The main filtering parameters were as follows: only biallelic SNPs, mean DP = 2.5, max-mean DP = 10, minDP = 2, max-missing = 0.5, IndelGap and SnpGap = 10 bp, and minor allele frequency ≤ 5%. We also excluded with low QUAL scores (QUAL < 20), did not allow monomorphic sites (only alternate allele), and removed sites with extremely high proportions of heterozygosity (≥ 75%), as we expect these to be either multiple-mapped sites not present in the genome assembly and/or the result of chimeric assembly of repetitive regions.

### Admixture and genomic PCA

We used several approaches to explore population structure across the hybrid zones. First, we used plink2 v.2.00a3.7 (Chang et al., 2015) on the “final filtered” .vcf to remove SNPs with r2 > 0.1 in 50 SNP windows, sliding every 10 SNPs along the genome. This resulted in 893,961 SNPs (Table S3). This output was then used to estimate population structure in Admixture v.1.3.0 (Alexander et al., 2009), with default parameters, and 5-fold cross-validation for K = 1–5. We also generated a genomic PCA in plink2, using the same input as used for Admixture. Additionally, we estimated the elevation for each point using the R package *elevatr* v.0.99.0 (Hollister, 2017), and conducted an ANOVA with a chi-squared test to determine whether elevation significantly explained variation in admixture proportion.

### Genotype imputation

We phased and imputed genotypes in BEAGLE v.5.4 (Browning et al., 2021). Our input .vcf file for genotype phasing and imputation was the filtered version as described in the variant filtering section (7,322,630 SNPs; Table S3), except we additionally only allowed sites with ≤ 5% missing data. Imputation was performed with default settings. Genotype imputation is expected to be more accurate when a large, independent reference panel is available. Although no reference panel is available for our study, recent research suggests that for suitably large data sets (several hundred samples or more), accurate imputation can still be achieved for low coverage WGS data even in the absence of a reference panel (Topaloudis et al., 2026). Furthermore, many downstream applications (including GWAS) are reportedly robust to imputation without a reference panel, though caution is advised for some inferences such as the comparison of genomic regions identified as homozygous-by-descent (Topaloudis et al., 2026).

### Local ancestry inference

Local ancestry inference was performed in ancestryHMM v.1.0.2 (Corbett-Detig & Nielsen, 2017). Samples with >99.999% assignment probability to either population at K = 2 in the Admixture analysis were designated as “pure” parent samples. This resulted in 47 *P. davidsonii* and 27 *P. newberryi* samples. Using the imputed genotype data set, we found SNPs with an allele frequency difference between parent species ≥ 0.75. We then LD-pruned this set of SNPs in 50bp windows, sliding 10bp, with r2 > 0.2 in plink2. This LD-pruning approach in each parent population separately resulted in 44,095 SNPs (Table S3) for the analysis and should reduce the effect of ancestry-specific linkage-disequilibrium. The nonoverlapping sets of SNPs from these analyses were then converted from .vcf into ancestryHMM format using a modified version of the vcf2ahmm.py script provided by the developers. We then ran ancestryHMM on admixed (not “pure” parent) individuals, fitting a model with two populations and using the global ancestry proportions inferred from Admixture as fixed variables for each population. This model assumed an initial population of *P. newberryi*, and a subsequent pulse of *P. davidsonii* ancestry replacing a proportion of individuals equal to the *P. davidsonii* global ancestry proportion. The timing of both ancestry pulses (initial *P. newberryi* population and subsequent *P. davidsonii* admixture) were both optimized with initial estimates at 100 generations, and the error rate per site was set to 1×10-3. For each SNP in each sample, ancestry was assigned if the posterior probability for homozygous *P. newberryi* (*new*), heterozygous (*het*), or homozygous *P. davidsonii* (*dav*) ancestry was ≥ 0.8. The average ancestry proportion across all sites was subsequently calculated to generate a genome-wide ancestry proportion for each admixed sample. We additionally calculated hybrid index and interclass heterozygosity for each admixed individual from these ancestry proportions, where hybrid index (*HI*) = *dav* + *het*/2, and interclass heterozygosity = *het*. These values were then plotted as a hybrid index-by-heterozygosity triangle plot (Figure 2f) with the theoretical maximum-heterozygosity curve under Hardy-Weinberg equilibrium (heterozygosity = 2*HI* * (1–*HI*)) shown for reference.

### Genomic landscape of divergence

Parental samples were identified as described in the local ancestry inference section. To calculate divergence and differentiation metrics for parent samples, we re-called variants for all parent samples using the same filters as described previously, except we additionally retained invariant sites (“all-sites” VCF file). This all-sites file, which contained 8,189,072 SNPs (Table S3), was used as input for pixy v.1.2.7.beta1 (Korunes & Samuk, 2021), in which we estimated FST, π, and dxy in 100kb non-overlapping windows across the entire genome, and in 100bp non-overlapping windows spanning the floral hue association peak. To examine relationships among metrics of diversity and differentiation, we estimated Pearson’s correlation coefficient between pairs of these statistics. These tests were performed across 100kb genomic windows. We calculated average π as the average value of π for *P. davidsonii* and *P. newberryi* in each window, following Shang et al., (2023).

### Floral trait differentiation

We employed a Random Forest classifier with 500 decision trees using the R package *randomForest* v.4.7-1.2 (Breiman, 2001) to identify traits most strongly differentiating parent species using all phenotypic measurements, including spectral reflectance data. Prior to the RF classifier, all missing data were imputed using the R package *mice* v.3.17.0 (Van Buuren & Groothuis-Oudshoorn, 2006). Any traits with > 3% relative importance, defined as the proportion of mean decrease in Gini for each trait divided by the sum of the mean decreases in Gini across all traits (see Figure S6), were included in downstream analyses.

To assess floral trait differentiation including hybrids, we first generated correlation matrices between traits, after removing one trait (crease to top height) due to its strong correlation (*r* = 0.95) with another trait (tube height at mouth) for downstream analyses. We then performed a PCA with floral traits, first imputing missing values for hybrids in the R package *mice*, using the pmm method, and scaling data prior to ordination. We then used generalized linear models (GLMs) with binomial error distributions and logit link functions to test whether PC1 predicted K1 and elevation. Model significance was assessed using a likelihood ratio test (LRT), and model fit was quantified using Nagelkerke’s pseudo-R2.

### GWAS

We performed admixture mapping on hybrid individuals that display continuous variation in traits due to recombination of parent species’ genotypes. This approach allowed us to identify genomic regions associated with trait variation. For modified hue and chroma associations, n = 283. For all other analyses, n = 282 (one individual for which sequencing and reflectance spectroscopy data were collected was inadvertently not photographed and phenotyped). To assess whether phenotypic traits conformed to assumptions of normality, we applied the Shapiro-Wilk test to each trait. Traits with p-values greater than 0.05 were considered approximately normal and retained without modification. For traits deviating from normality, we evaluated common transformations, including logarithmic (for strictly positive values), square-root (for non-negative values), and Box-Cox transformations. For each transformation, we repeated the Shapiro-Wilk test and selected the transformation yielding the highest p-value, provided it improved normality relative to the untransformed data. If no transformation improved normality, the trait was retained in its original form. A complete list of p-values and trait transformations are provided in Table S10.

The imputed SNP data set was LD-pruned using the same parameters applied in the Admixture analysis, removing SNPs with r2 > 0.1 in 50 SNP windows and sliding every 10 SNPs along the genome. The resulting LD-pruned SNP set (344,164 SNPs; Table S3) was used to calculate the centered relatedness matrix in GEMMA v.0.98.5 (Zhou & Stephens, 2012). This relatedness matrix, together with the full imputed SNP data set (3,021,696 SNPs; Table S3), was used to perform GWAS under the univariate linear mixed model (ULMM) implemented in GEMMA. The inclusion of the relatedness matrix as a random effect in the mixed model helps account for confounding effects of population structure (Zhou & Stephens, 2012). To determine appropriate significance thresholds, we applied two approaches. First, we used a Bonferroni correction at α = 0.05 across the full imputed SNP set. Second, we implemented a permutation-based approach, in which phenotypic values for each trait were randomly shuffled 100 times, and the ULMM was rerun for each permutation. From each run, we collected the lowest 1% of p-values, and the empirical threshold for each trait was set at the 0.0001% quantile of this pooled distribution. Both approaches yielded comparable thresholds; however, we report results using the permutation-derived thresholds, as they rely on fewer assumptions about the underlying distribution of test statistics.

We defined association peaks as clusters of at least five SNPs exceeding the genome-wide significance threshold. We identified protein-coding genes located within 20 kb upstream or downstream of each association peak using the *P. davidsonii* reference genome annotation, modified to retain only the longest isoform per gene model. This 20kb window yields an inclusive set of genes in the vicinity of each identified association peak (Hooper et al., 2024; Schield et al., 2024; Walsh et al., 2023). Candidate genes were then examined manually, with particular attention to those previously implicated in, or plausibly related to, the focal traits. Of species-diagnostic traits, only floral hue and tube height at the inflection point produced clear association peaks.

### Phylogenetic inference of candidate gene near top floral hue GWAS SNP

To infer the identity of the candidate gene located near the top floral hue GWAS SNP, we constructed a maximum-likelihood phylogeny using putative homologous protein sequences from multiple angiosperm lineages. We assembled three sets of reference sequences: (1) canonical dihydroflavonol-4-reductase (DFR), (2) canonical anthocyanidin reductase (ANR), and (3) the closest BLASTP matches to the unknown GWAS candidate gene. Protein sequences were obtained from *Arabidopsis thaliana* and *Glycine max*, several *Penstemon* species (*P. davidsonii*, *P. barbatus*, *P. eatonii*, *P. smallii*, and *P. neomexicanus*) and several Lamiales species (*Paulownia fortunei*, *Buddleja alternifolia*, *Rehmannia glutinosa*, *Sesamum alatum*, *Orobanche gracilis*, *Perilla frutescens* var. *hirtella*, and *Orobanche minor*). Only DFR was available for *P. neomexicanus*. Because a clear ANR ortholog could not be confidently identified in Lamiales species, only canonical ANR sequences from *A. thaliana* and *G. max* were included. As an outgroup, we incorporated cinnamoyl-CoA reductase (CCR) protein sequences from *A. thaliana* and *G. max*. The full list of protein sequences and corresponding GenBank accessions are provided in Table S11. Sequences were aligned in MUSCLE v2.8.1551 (Edgar, 2004) using default parameters, and alignment trimming was performed with trimAl v1.5.rev1 (Capella-Gutiérrez et al., 2009) using the automated1 setting. The phylogeny was estimated with IQ-TREE v2.3.6 (Minh et al., 2020) using ModelFinder to select the best-fit substitution model, and 1,000 UFBoot replicates and 1,000 bootstrap replicates for the Shimodaira-Hasegawa-like approximate likelihood ratio test (SH-aLRT) to assess branch support (Guindon et al., 2010).

### Floral pigment extraction

We determined the identity of anthocyanidin pigments produced in floral tissues, using dried floral tissues from the same flowers used for phenotyping. We used an isoamyl alcohol extraction protocol outlined in Harborne (1984) and separated pigments using TLC on Cellulose F glass plates (Millipore, Burlington, MA, USA) using a forestal solvent (glacial acetic acid:H2O:HCl 30:10:3). We compared extracted anthocyanidin pigments to a multi-anthocyanidin standard solution containing pelargonidin, cyanidin, and delphinidin each in concentrations of ∼0.013 mg/mL, as described in Stevens et al., (2023). Dried TLC plates were photographed in a controlled light box using standardized camera settings (Nikon z50 camera, shutter speed: 1/125, aperture: f16, iso: 500) and lighting conditions (550D LED ring light). Photographs were imported into the R package *qTLC* v.1.0 (Mac Fhionnlaoich et al., 2018) with the MinPeakProm parameter set to 6 and the divisor parameter set to 1.4. After spot intensities were calculated, values were averaged across replicates for each standard dilution prior to fixing a regression between values to infer the relationship between spot intensity and pigment quantity. For each pigment spot, intensity was converted to pigment quantity by comparing values to the standard curve. Pigment production for the entire flower was then estimated by accounting for the elution volume. Total pigment mass fraction in mg/g (mg pigment quantity) / (g dry flower mass) was then calculated by dividing total estimated pigment quantity by dried flower mass.

### Pollinator visual modeling

Parental and hybrid samples were identified as described in the Admixture analyses. We estimated the detectability of each floral spectrum under bee vs. bird visual models using *pavo* (Maia et al., 2019). The bee visual system was represented by a honeybee (*Apis mellifera*) visual model (Menzel & Blakers, 1976), and the hummingbird visual system was represented by the average avian violet-sensitive model, as North American hummingbirds are not predicted to be ultraviolet-sensitive, based on the opsin gene *SWS1* (Ödeen & Håstad, 2010). Spectral reflectance data were first clipped to 300-700 nm to match the visual sensitivity ranges of pollinators. For samples with multiple flowers, reflectance spectra were aggregated to produce a single representative spectrum per individual. We used the reflectance spectra of leaf tissue from *Penstemon x jonesii*, measured without using microscope slides, as a background for visual models (Stevens, 2025). We then used the function *vismodel* to calculate quantum catches for each photoreceptor for each visual system. Quantum catches were calculated using ideal lighting conditions, default quantum catch parameters (Qi), no von Kries transformation, and using the long wavelength receptor for each visual system to calculate achromatic stimulation. This output was then used with the function *coldist* to calculate distances between colors (and thus perceptual discriminability) in pollinator visual space using a receptor-noise model (Vorobyev & Osorio, 1998). This model expresses color contrast as just noticeable differences (JNDs), where values < 1 JND are considered perceptually indistinguishable, 1 JND corresponds to the approximate threshold for discrimination, and values > 1 JND indicate increasingly distinct colors. We used cone proportions of 1:2:1 (UVs, S, M) for the trichromatic bee visual system (Menzel & Blakers, 1976) and 1:1:1:2 (Vs, S, M, L) for the tetrachromatic avian visual system (Ödeen & Håstad, 2010). All other parameters were left at default values. We ran one-way ANOVAs and post-hoc Tukey’s HSD tests to test for statistical differences among sample classes (*P. davidsonii*, hybrid, and *P. newberryi*) within each visual model type. For the honeybee visual system, we used the function *colspace* to understand what photoreceptors are used to perceive various colors. To achieve this we used output from the *vismodel* function with hyperbolic-transformed quantum catches (Ei), the von Kries correction applied, and an average green foliage background.

### Ancestry at candidate loci for floral hue

To genotype samples at the *DFR-like* candidate gene, the 10 significant GWAS SNPs within the gene model were extracted from the imputed SNPs .vcf file for all hybrids. All SNPs were either fixed between parent species or contained a single discordant allele between parent species (Table S12). Because these SNPs were ancestry informative, we used them to assign locus-level ancestry at the *DFR-like* candidate gene. For each sample, genotype was assigned at each SNP, and locus-level ancestry was determined by the percentage of SNP genotype calls, such that ≥50% heterozygous was classified as heterozygous ancestry, and >50% for either parent genotype was classified as homozygous for that parental ancestry. A paucity of ancestry-informative SNPs in the *F3’5’H* gene model led us to genotype at this candidate gene using PCR, rather than with a bioinformatic approach. To genotype samples at *F3’5’H*, we generated primers and ran a PCR. Primer sequence and PCR conditions are described in Table S13. The *P. newberryi* ancestry at this gene contains a large deletion which encompasses the entire second exon and causes loss-of-function (Stone & Wessinger, 2024), enabling genotyping by visualizing *F3’5’H* genotype by eye. We then fit three nested models to estimate the role that ancestry at *DFR-like* and *F3’5’H* may play in affecting flower color. Our models were as follows: model 1: main effect of *F3’5’H* only; model 2: additive effects of *F3’5’H* and *DFR-like*; model 3: full interaction model, including main effects of *F3’5’H* and *DFR-like*, and their interaction. We then compared models with an ANOVA and assessed model fit with AIC.

### Genomic Clines

We used hierarchical Bayesian genomic cline analysis to estimate locus-specific patterns of introgression relative to the genome-wide admixture gradient, as implemented in *bgchm* (Gompert et al., 2024). This method employs the logit-logistic cline function proposed by Fitzpatrick (2013), where the probability that an allele at locus *i* in individual *j* was inherited from population 1 is given by:

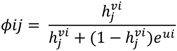

with *h* representing the proportion of the genome inherited from population 2 (hybrid index), *v* describing the cline slope relative to the genome-wide average (*v* = 1), and *u* relating to the cline center. Following Bailey (2024), we expressed the cline center parameter as:

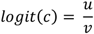

such that *c* denotes the hybrid index at which *ϕ* = 0.5. We measured variance in clines as the variance in log(*v*) and logit(*c*), which each have expected means of 0.

To estimate genomic clines, we first identified all SNPs within floral trait GWAS association peaks with minor allele frequency (MAF) > 0.1. Floral trait loci were defined as all gene models within 20kb of any SNP exceeding the permutation-derived genome-wide significance threshold in the GWAS. Because pseudochromosomes 1086 (containing the major floral hue locus) and 2532 (associated with lower petal angle) contained disproportionately large numbers of significant SNPs, we randomly subsampled 10% of SNPs from each, while retaining all other significant SNPs from other pseudochromosomes. We next selected background SNPs (MAF ≥ 0.1) located > 20kb from any significant SNP and outside all floral trait loci. From these, 2% of SNPs per pseudochromosome were randomly sampled. In total, genomic clines were estimated for 49,019 SNPs.

Hybrid indexes were estimated for each individual using 1,000 randomly selected SNPs from the same subsampling scheme. The cline standard deviation parameters were calculated from this subset and used as point estimates for all other cline analyses in *bgchm*. Genomic clines were estimated using the est_genocl command, with 4,000 iterations and a 2,000-iteration burn-in. Convergence was assessed in *R* using effective sample size (ESS) and *R̂* statistics. We considered cline gradient estimates as outliers if their 95% confidence intervals fell above 1 (for steep clines) or below 1 (for shallow clines). The distributions of cline slopes and centers in specific genomic regions (e.g., the hue GWAS association peak) were statistically compared with a Mann-Whitney U test.

### Excess heterozygosity

To quantify excess heterozygosity across the genome, we first filtered the dataset to include only hybrid individuals with admixture proportions between 25–75% and only SNPs fixed between parent species in the imputed .vcf (136,590 SNPs; Table S3). For these sites, observed and expected genotype counts were calculated using vcftools with the --hardy function, which computes Hardy–Weinberg expected genotype frequencies based on allele frequencies estimated from the hybrid subset. From these counts, we obtained the proportion of observed heterozygotes and the expected proportion under Hardy–Weinberg equilibrium at each site, and defined heterozygosity excess as *Hexcess = P(Ho) – P(He)*.

### Data Availability

Raw sequencing reads for all samples used in this study are available at NCBI BioProject PRJNA1403563. Other data underlying this study are available at: https://datadryad.org/share/LINK_NOT_FOR_PUBLICATION/jnD7cENkYCMMNQxBQE4PXg9i5heLVD4RIdecXYP6el8. All code necessary for this study is available at https://github.com/benstemon/sierra-hybrid-zone.

## Data Availability

Raw sequencing reads for all samples used in this study are available at NCBI BioProject PRJNA1403563. All code necessary for this study is available at https://github.com/benstemon/sierra-hybrid-zone.

## Author Contributions

B.W.S. and C.A.W. conceived the project. B.W.S., N.H.W., T.H.D., Z.J.R., and A.M.M. collected the data. B.W.S. analyzed the data and generated figures. B.W.S. wrote and revised the manuscript with contributions from C.A.W.

## Acknowledgments

We thank the USFS for sampling permits at both hybrid zones. We gratefully acknowledge the computational resources provided by the Hyperion high performance computing cluster at the University of South Carolina, and the technical assistance and resources provided by Research Computing at the University of South Carolina (RRID:SCR_027488). We also thank M.L. Smith and two anonymous reviewers for helpful comments on the manuscript, A. Childs for assistance with pigment concentration analysis, and M. Stewart for assistance with PCR primer development.

