## Supplemental Materials for "Genomic signatures of reproductive isolation are decoupled from floral divergence in a long-standing hybrid zone"

### Supplemental Results

#### *Comparison of reference bias*

We observed small, consistent differences in mapping performance depending on reference genome. Individuals of each parental species (determined by Admixture results: see main text) had slightly higher coverage and depth, and less missing data, when mapped to its own reference (Figure S1). Hybrid individuals exhibited an intermediate pattern with a slight shift toward the *P. newberryi* reference. Despite these differences, the total effect was small for both depth (mean differences: *P. davidsonii* $\approx$ 0.21x $\pm$ 0.06x, hybrids $\approx$ 0.15x $\pm$ 0.16x, *P. newberryi* $\approx$ 0.30x $\pm$ 0.04x) and missingness (mean differences: *P. davidsonii* $\approx$ 2.9% $\pm$ 0.2%, hybrids $\approx$ 2.3% $\pm$ 2.5%, *P. newberryi* $\approx$ 4.6% $\pm$ 0.4%). Summary statistics for these metrics are provided in Table S1. Importantly, the distribution for hybrids encompasses 0 (no net change), indicating that irrespective of the reference used, there would be some degree of mapping bias across hybrid samples. However, the lack of strong systematic bias in mapping rates indicates there should not be a substantial loss of data for either species. In light of this, and given the availability of a high-quality annotation for the *P. davidsonii* reference and a lack thereof for the *P. newberryi* reference, we opted to conduct all downstream analyses on the *P. davidsonii* reference genome.

### Supplemental Figures

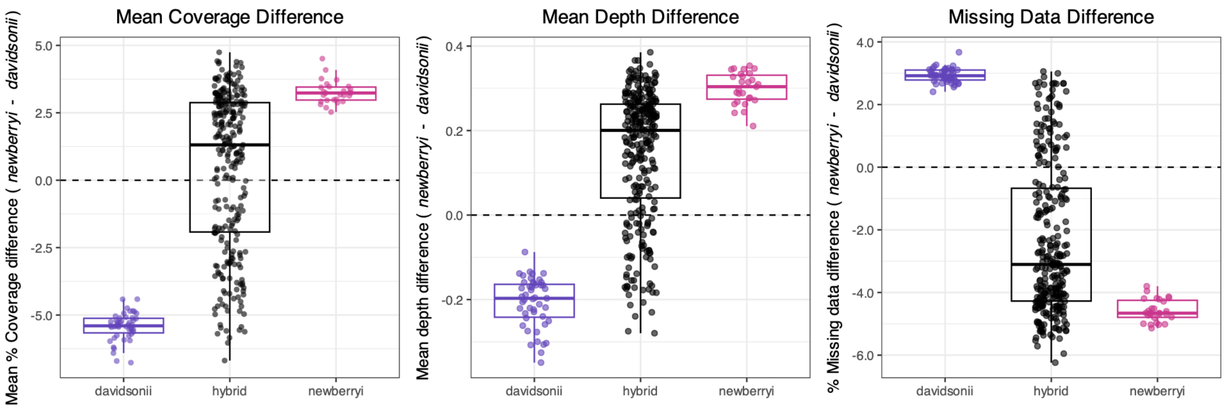


**Figure S1.** Comparisons of relative genome-wide coverage (left), mapping depth (center), and missingness (right) when mapped to the *P. newberryi* reference genome vs. the *P. davidsonii* reference genome. Individuals were assigned to "pure" *P. davidsonii*, hybrid, or "pure" *P. newberryi* based on Admixture results. In each plot, a value of 0 would indicate no difference for that individual when mapped to each reference.

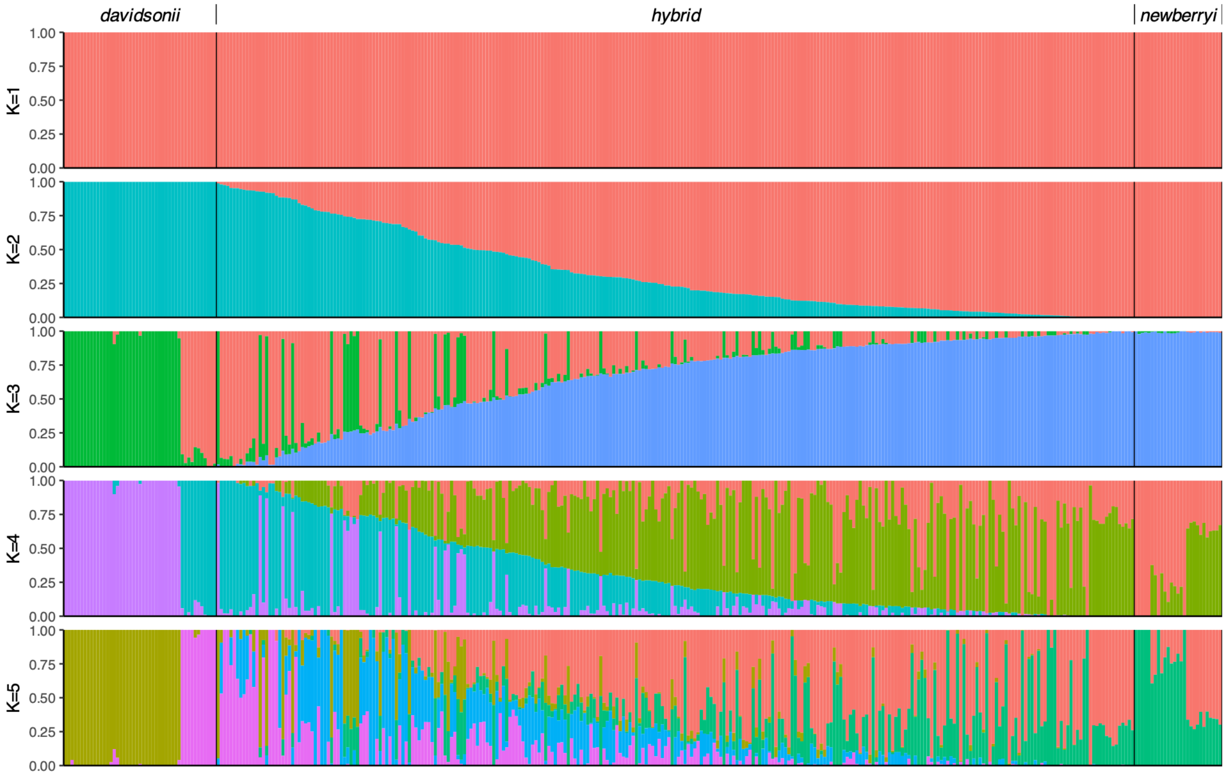


**Figure S2.** Bar plots of the Admixture analysis, from K = 1–5. Each column is an individual and each row represents a different value of K; individuals are in the same order across rows. Individuals are labeled “davidsonii”, “hybrid”, or “newberryi” based on their assignment from K = 2.

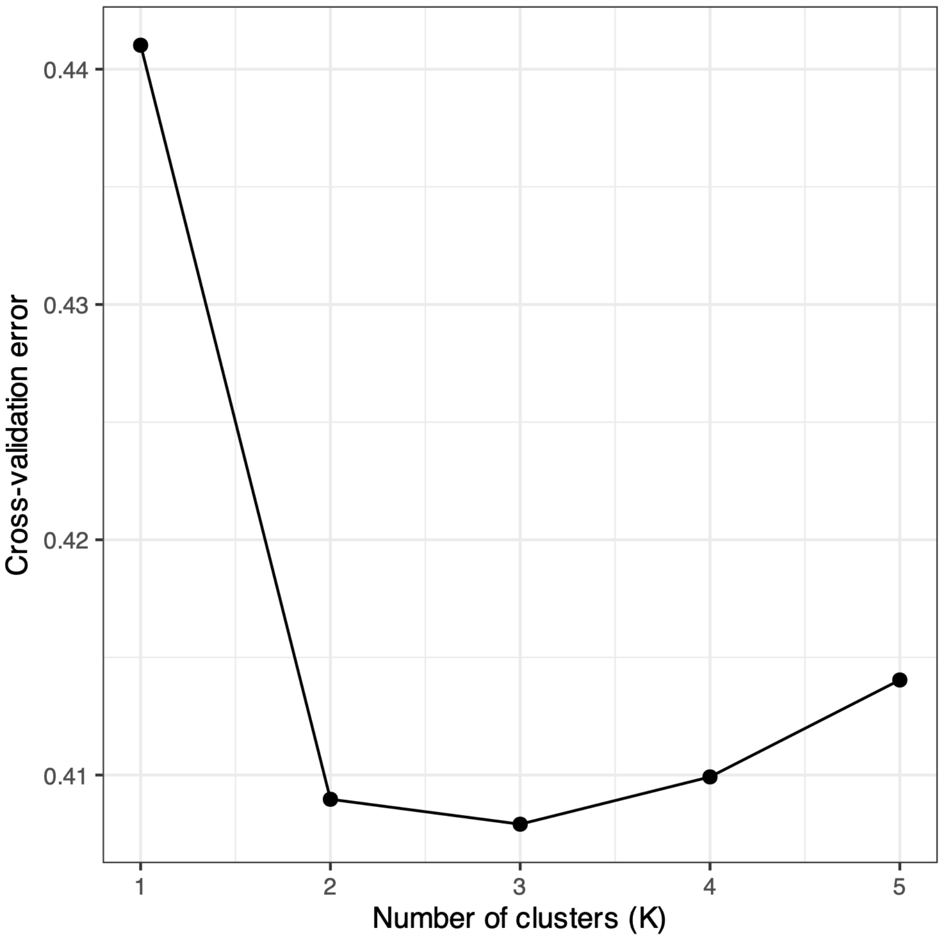


**Figure S3.** Cross-validation error of the Admixture analysis for K = 1–5.

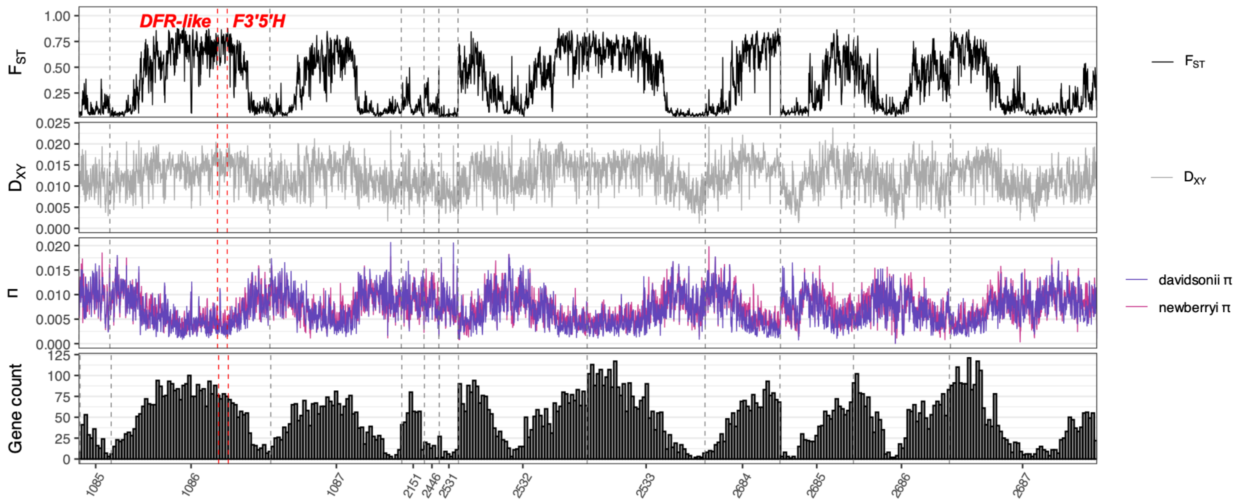


**Figure S4.** Genome-wide patterns of $F_{ST}$, $d_{xy}$, nucleotide diversity (π), and gene count in 100kb non-overlapping windows between "pure" *P. davidsonii* and *P. newberryi*. Gene count histograms are based on annotations for the *P. davidsonii* reference genome.

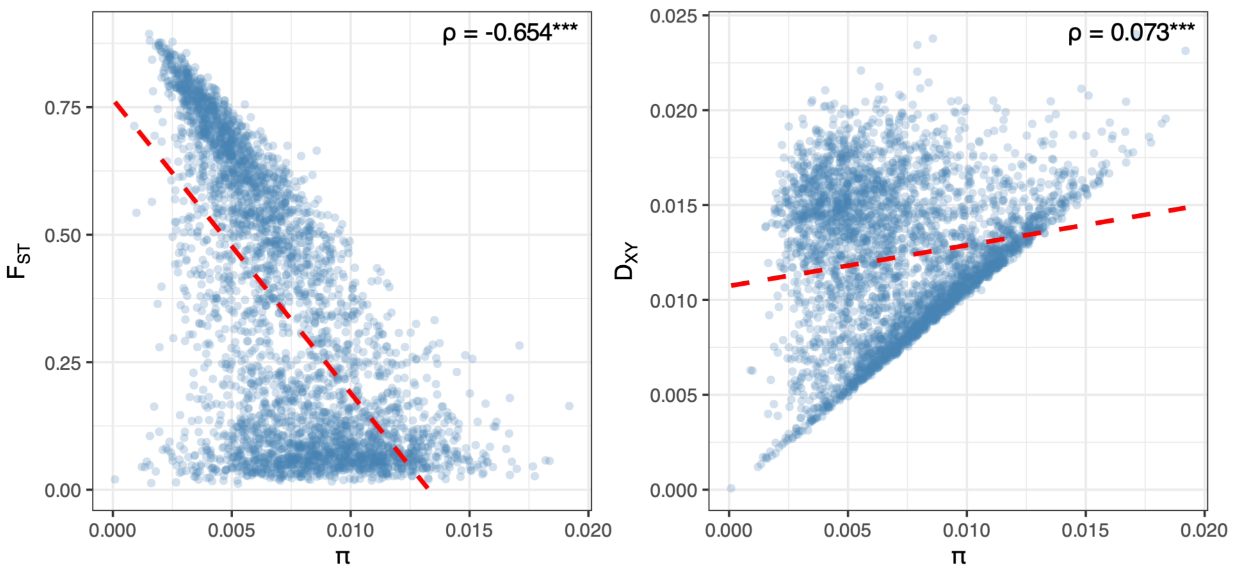


**Figure S5.** Genome-wide correlation analysis for π vs. $F_{ST}$ (left) and π vs. $d_{xy}$ (right) between parent species in 100kb windows. Pearson's ρ with $p < 0.001$ are denoted with three asterisks.

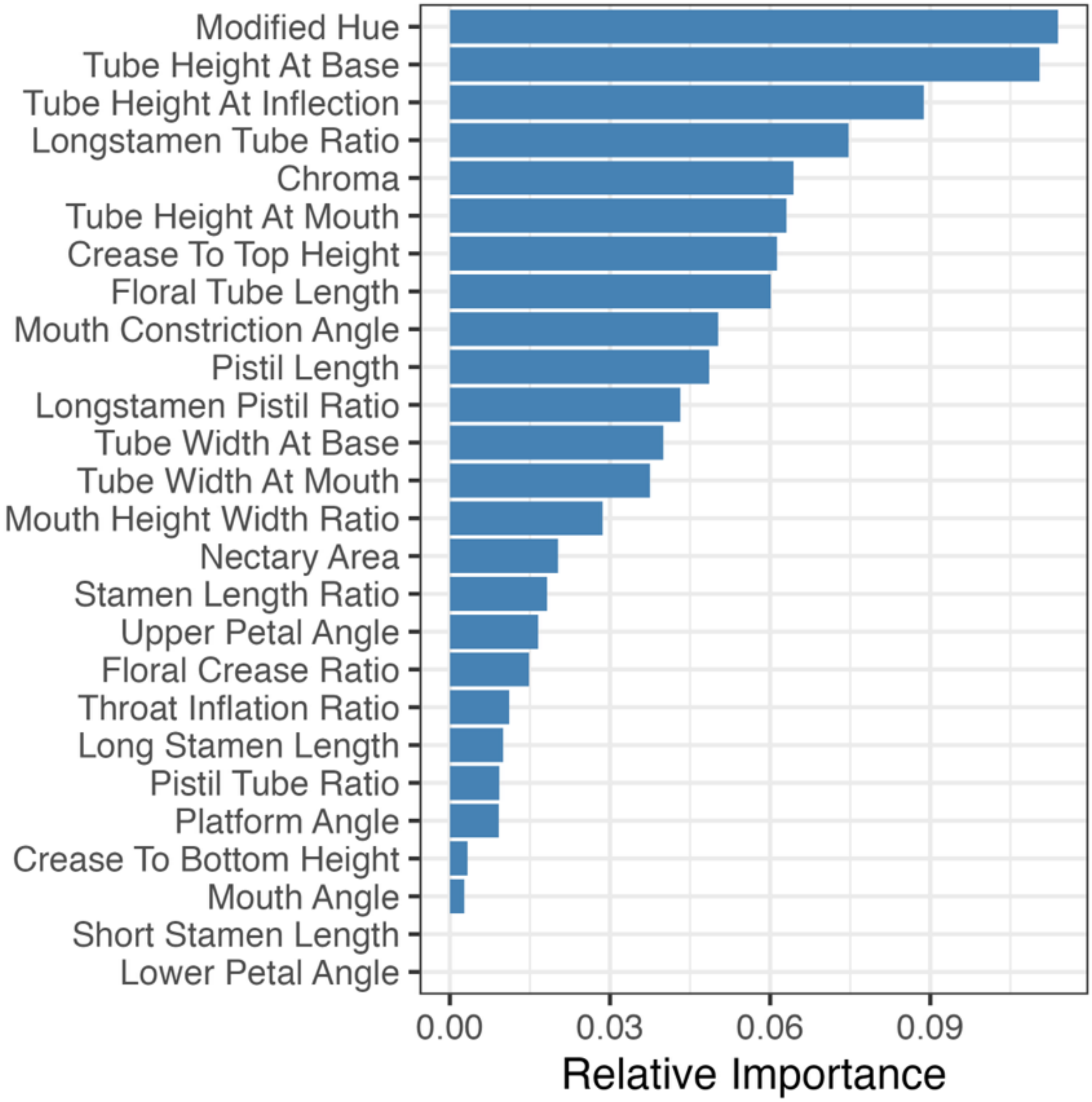


**Figure S6.** Relative importance metrics as identified by the Random Forest classifier. Relative importance was calculated as the proportion of mean decrease in Gini for each trait divided by the sum of the mean decreases in Gini across all traits.

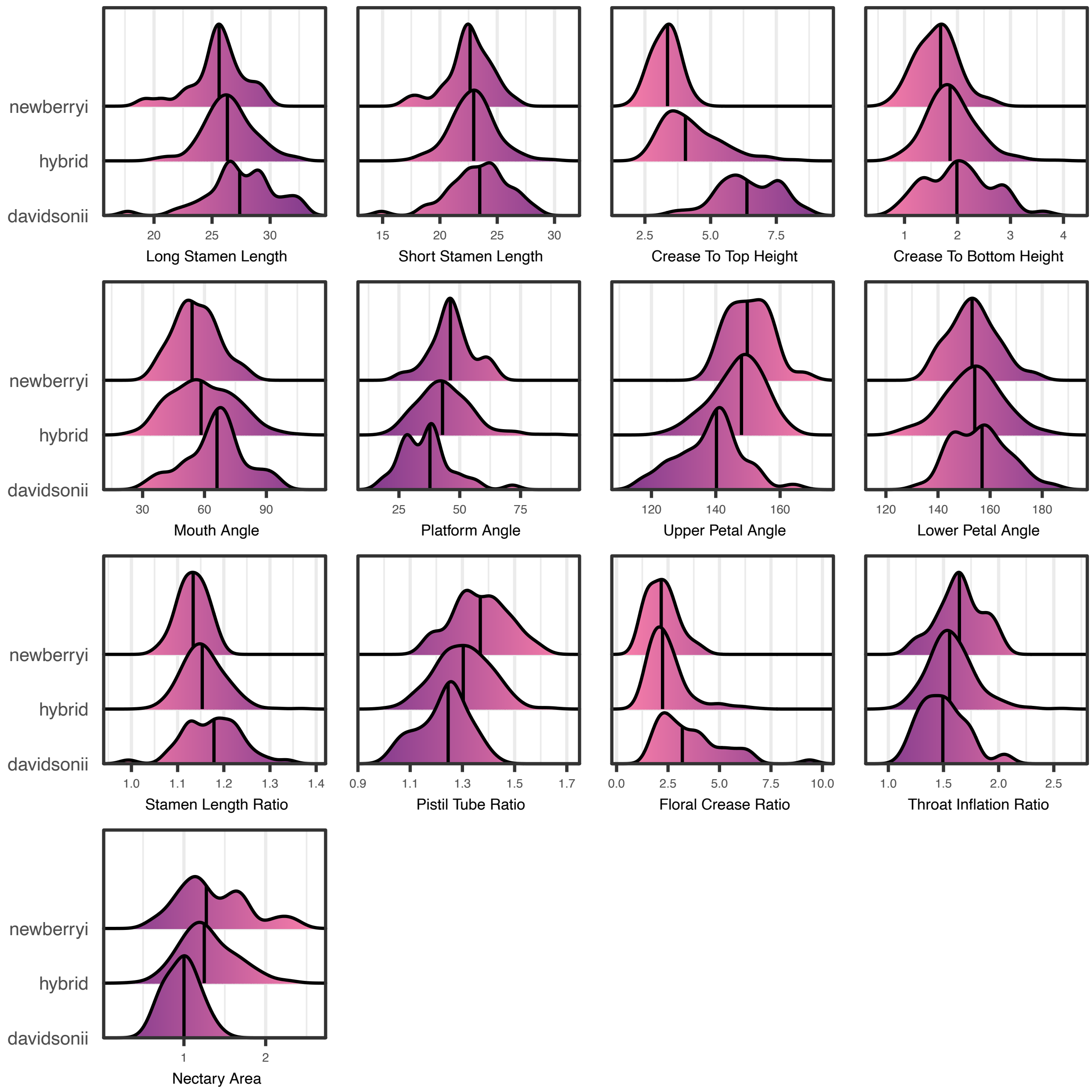


**Figure S7.** Ridgeline plots of all traits not identified as "important" in the Random Forest classifier for parental and hybrid individuals.

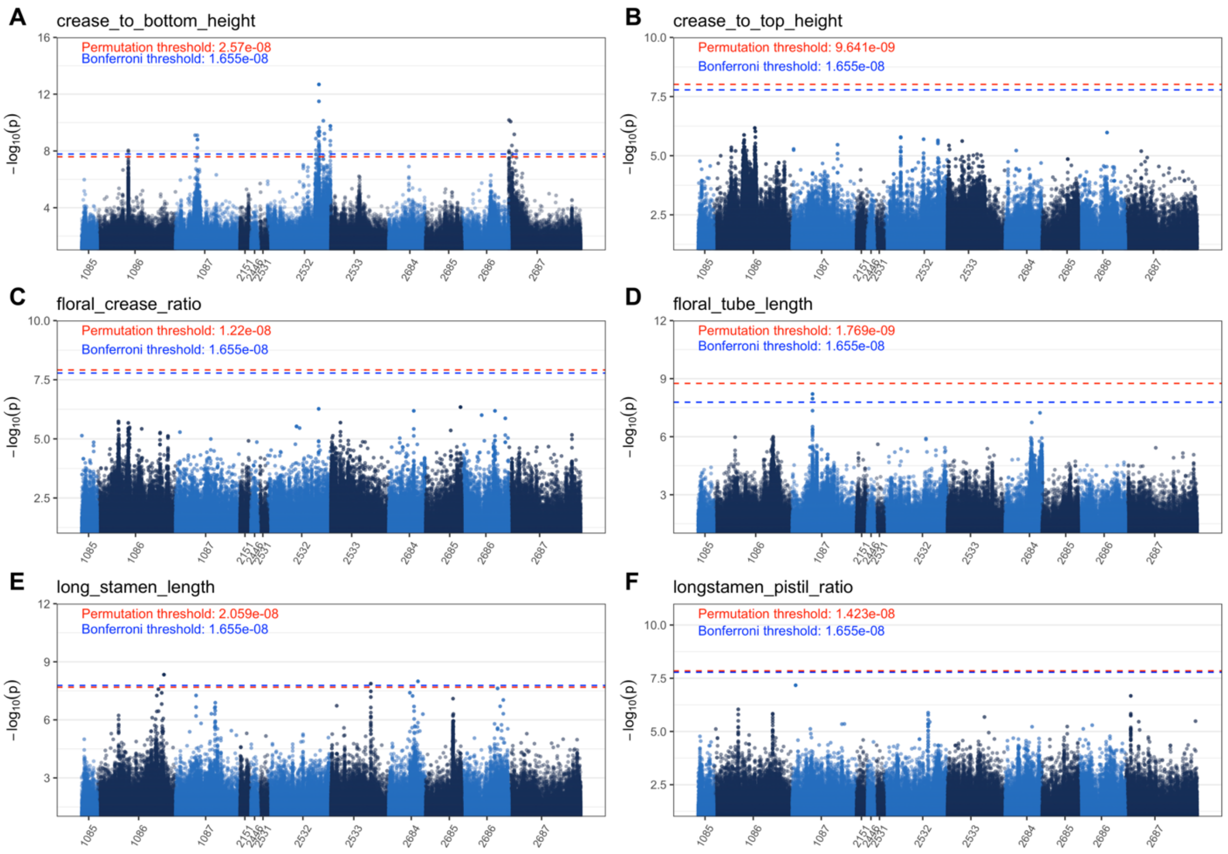


**Figure S8**. Manhattan plots of GWAS results for six traits. Panels A-F correspond to crease to bottom height, crease to top height, floral crease ratio, floral tube length, long stamen length, and long stamen to pistil ratio, respectively. The dashed red line denotes the genome-wide significance threshold as determined by permutation, and the blue dashed line denotes the Bonferroni-corrected significance threshold.

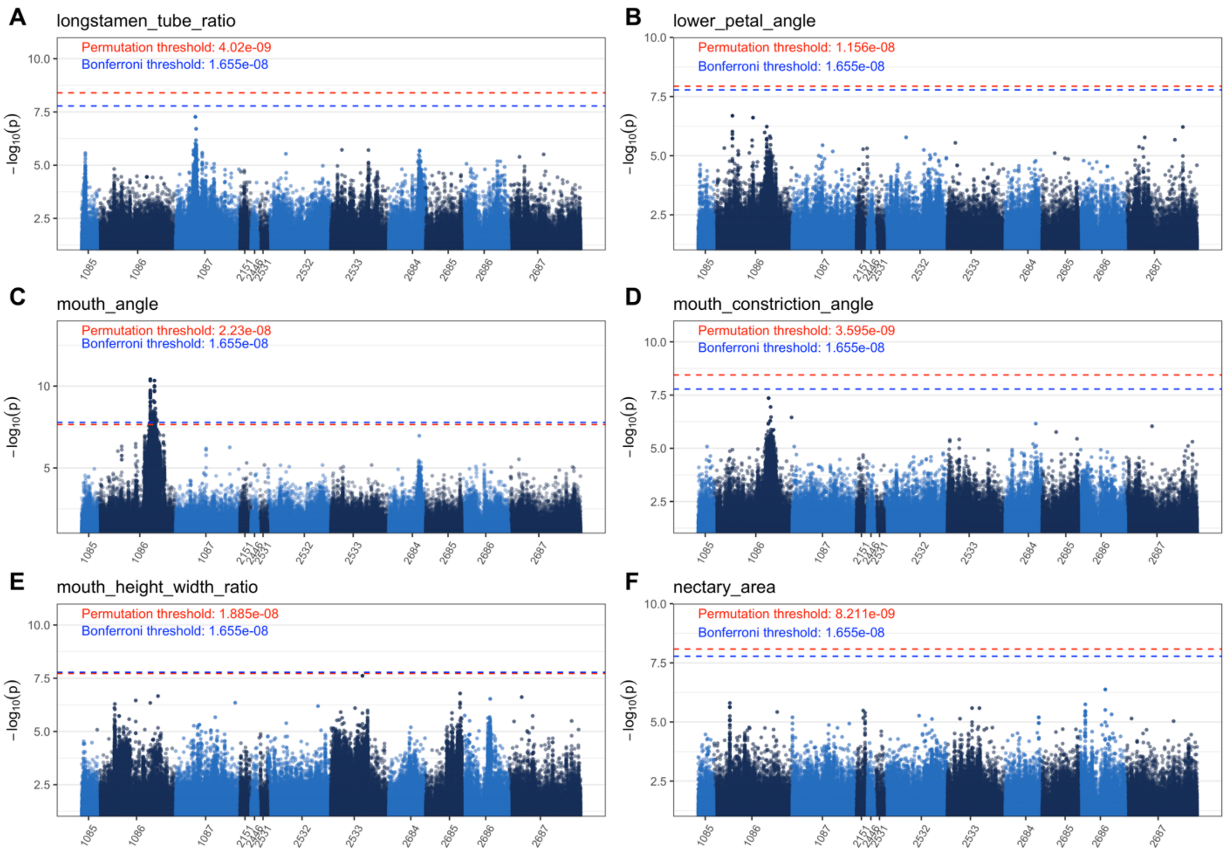


**Figure S9**. Manhattan plots of GWAS results for six traits. Panels A-F correspond to long stamen to floral tube ratio, lower petal angle, mouth angle, mouth constriction angle, mouth height to width ratio, and nectary area, respectively. The dashed red line denotes the genome-wide significance threshold as determined by permutation, and the blue dashed line denotes the Bonferroni-corrected significance threshold.

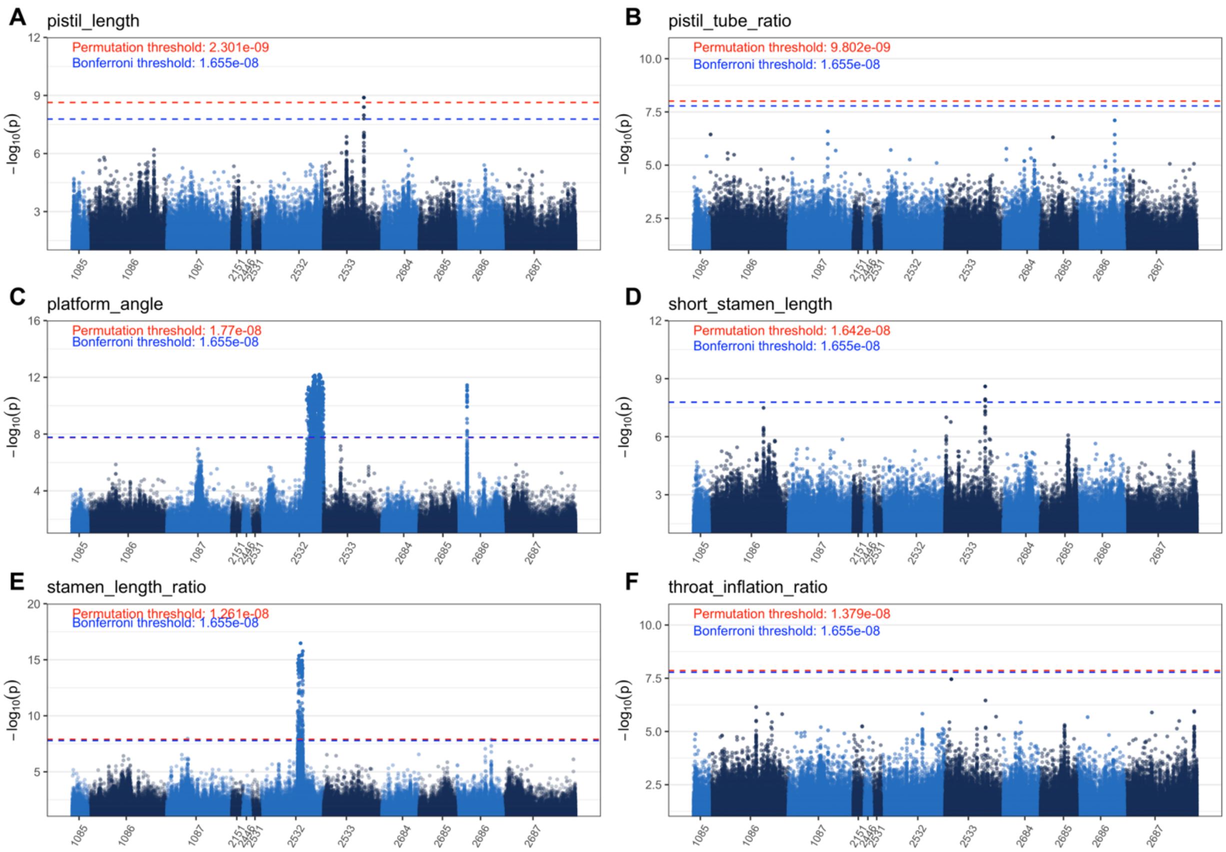


**Figure S10**. Manhattan plots of GWAS results for six traits. Panels A-F correspond to pistil length, pistil to floral tube length ratio, platform angle, short stamen length, stamen length ratio, and throat inflation ratio, respectively. The dashed red line denotes the genome-wide significance threshold as determined by permutation, and the blue dashed line denotes the Bonferroni-corrected significance threshold.

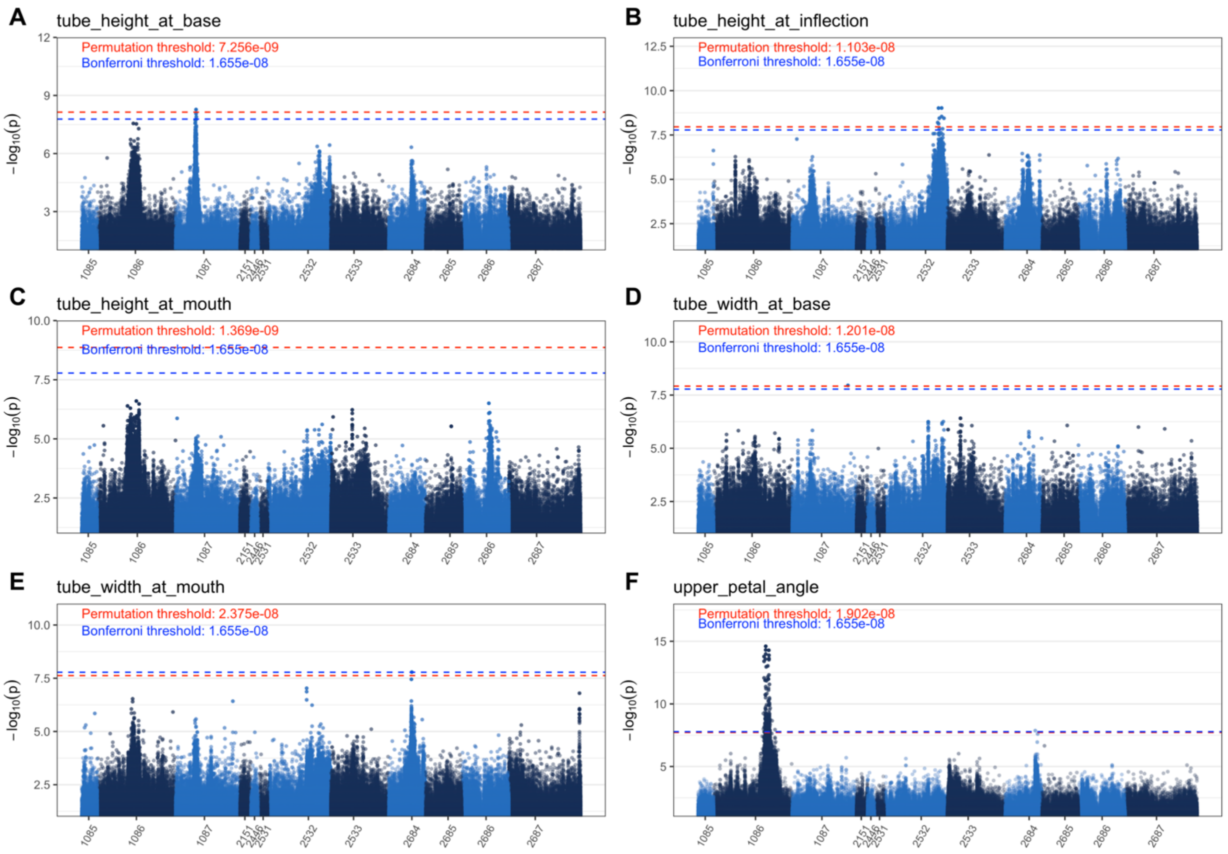


**Figure S11**. Manhattan plots of GWAS results for six traits. Panels A-F correspond to tube height at base, tube height at inflection, tube height at mouth, tube width at base, tube width at mouth, and upper petal angle, respectively. The dashed red line denotes the genome-wide significance threshold as determined by permutation, and the blue dashed line denotes the Bonferroni-corrected significance threshold.

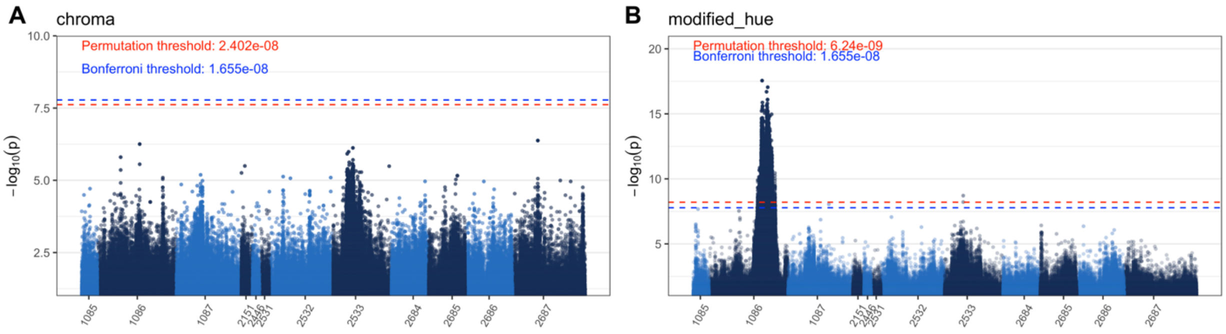


**Figure S12**. Manhattan plots of GWAS results for two traits. Panels A-B correspond to chroma and modified hue, respectively. The dashed red line denotes the genome-wide significance threshold as determined by permutation, and the blue dashed line denotes the Bonferroni-corrected significance threshold.

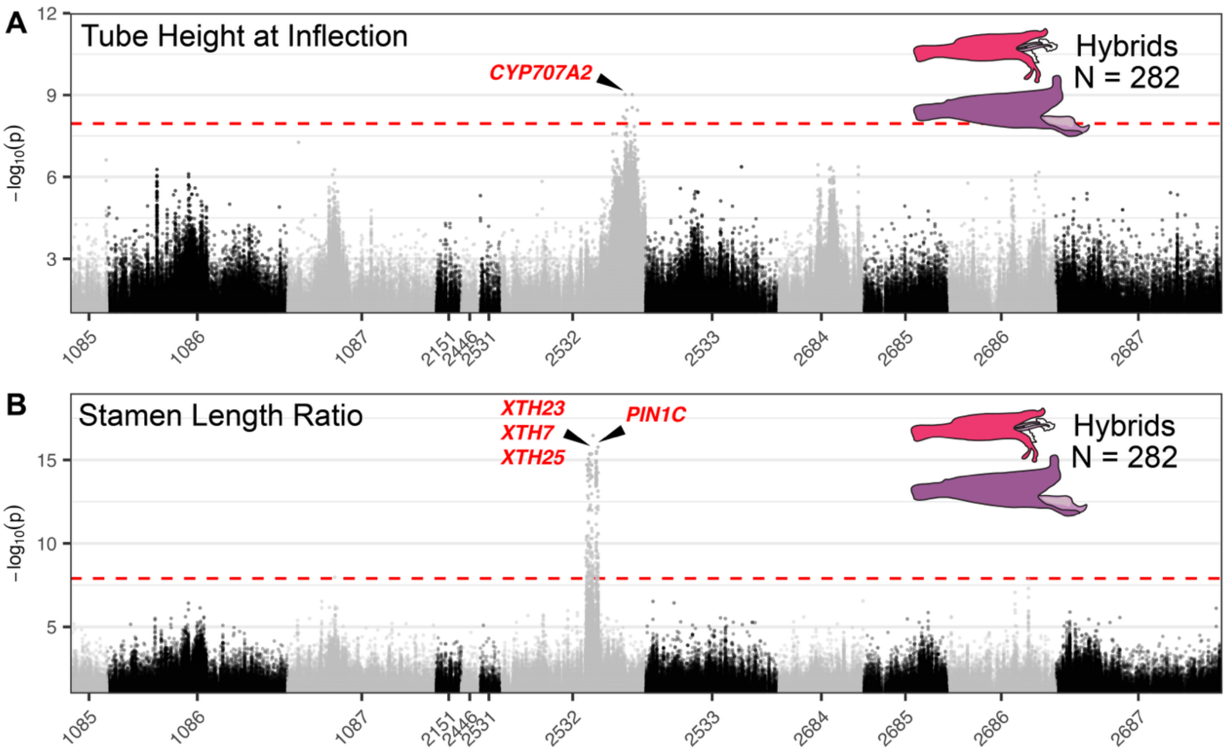


**Figure S13.** Annotated Manhattan plots of GWAS results for two traits. (A) GWAS of stamen length ratio. The Manhattan plot identified one large association peak on pseudochromosome 2532. The location of the candidate gene *CYP707A2* is highlighted. (B) GWAS of tube height at inflection. The Manhattan plot identified one large association peak on pseudochromosome 2532. For both plots, association support is plotted as -$\log_{10}(pwald)$, the dashed red line denotes the genome-wide significance threshold as determined by permutation, and the approximate locations of candidate genes of interest are annotated.

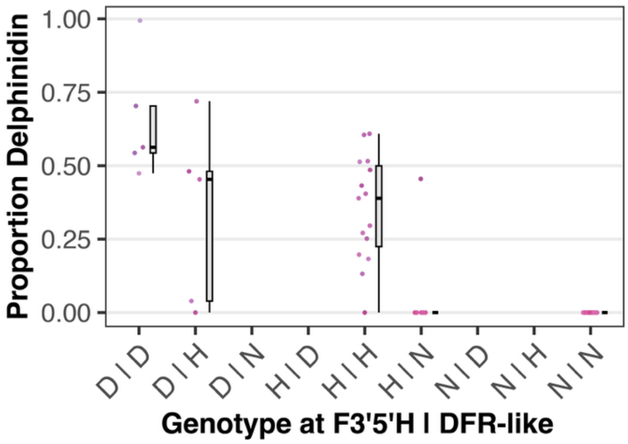


**Figure S14.** Values of delphinidin proportion with respect to genotype at *F3'5'H* and the *ANR/DFR* candidate genes within the hue GWAS association peak.

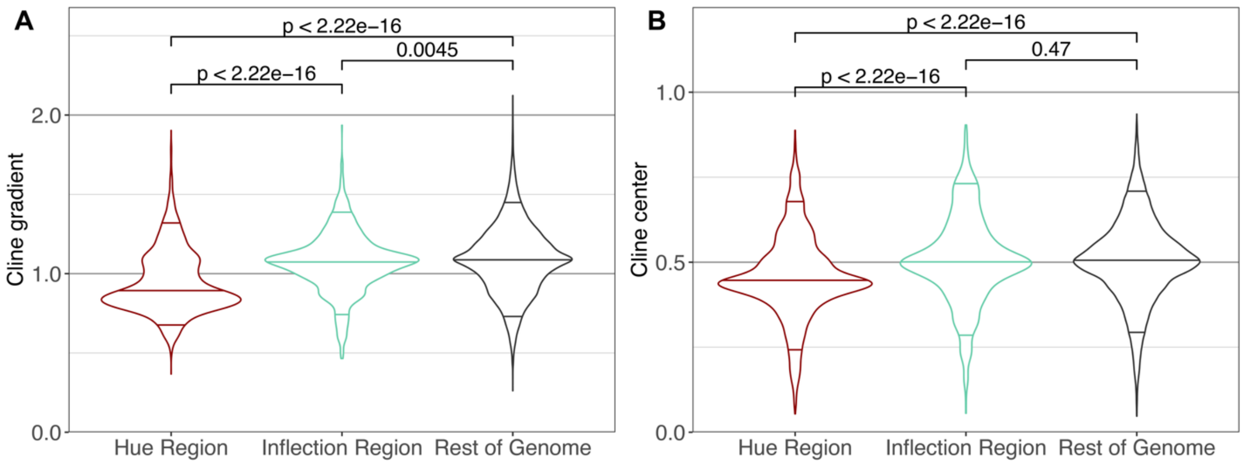


**Figure S15.** Distributions of cline gradients (A) and centers (B) for SNPs within the floral hue GWAS association peak, the tube height at inflection GWAS association peak, and the rest of the genome. Distributions were compared using Mann-Whitney U tests.

### Supplemental Tables

*Table S1. Mapping bias statistics per sample class*

| Class | n | Mean Coverage Difference | SD | Min | Max |
|---|---|---|---|---|---|
| *davidsonii* | 47 | -5.430 | 0.507 | -6.765 | -4.412 |
| hybrid | 283 | 0.436 | 2.903 | -6.688 | 4.744 |
| *newberryi* | 27 | 3.260 | 0.424 | 2.529 | 4.509 |
| **Class** | **n** | **Mean Depth Difference** | **SD** | **Min** | **Max** |
| *davidsonii* | 47 | -0.207 | 0.059 | -0.349 | -0.088 |
| hybrid | 283 | 0.145 | 0.156 | -0.280 | 0.385 |
| *newberryi* | 27 | 0.300 | 0.038 | 0.211 | 0.354 |
| **Class** | **n** | **Mean Missing Difference** | **SD** | **Min** | **Max** |
| *davidsonii* | 47 | 2.93% | 0.22% | 2.41% | 3.67% |
| hybrid | 283 | -2.34% | 2.41% | -6.24% | 3.06% |
| *newberryi* | 27 | -4.58% | 0.36% | -5.14% | -3.80% |

*Differences are calculated as the metric obtained after mapping to the P. newberryi reference minus the metric obtained after mapping to the P. davidsonii reference. Positive values indicate higher values when mapped to the P. newberryi reference.*

*Table S2. Read coverage and depth per scaffold (pseudochromosome)*

| Pseudo-chromosome | **1086** | **2687** | **1087** | **2532** | **2533** | **2686** | **2684** | **2685** | 1085 | 2151 | 2531 | 2446 |
|---|---|---|---|---|---|---|---|---|---|---|---|---|
| **Depth** | | | | | | | | | | | | |
| Min | 2.33 | 2.19 | 2.15 | 2.26 | 2.51 | 2.03 | 2.42 | 2.33 | 1.99 | 1.79 | 6.3 | 2.19 |
| Max | 6.82 | 6.37 | 6.31 | 6.33 | 7.29 | 5.88 | 6.9 | 6.77 | 5.81 | 5.14 | 18.4 | 6.75 |
| Mean | 4.97 | 4.67 | 4.57 | 4.69 | 5.37 | 4.44 | 5.15 | 5.01 | 4.36 | 3.71 | 12.06 | 4.92 |
| **Coverage** | | | | | | | | | | | | |
| Min | 68.9% | 62.0% | 61.3% | 64.9% | 65.9% | 60.4% | 67.2% | 61.3% | 55.8% | 57.5% | 50.4% | 54.8% |
| Max | 84.3% | 77.5% | 75.8% | 80.3% | 81.9% | 79.0% | 84.7% | 82.0% | 73.7% | 73.8% | 73.4% | 74.3% |
| Mean | 79.3% | 72.4% | 72.1% | 74.1% | 76.5% | 71.3% | 77.9% | 72.3% | 66.7% | 68.2% | 60.8% | 66.1% |

*Highlighted pseudochromosomes comprise the eight largest scaffolds.*

*Table S3. SNP counts*

| Analysis | SNPs | Info |
|---|---|---|
| Raw unfiltered .vcf | 56279850 | See main text |
| Filtered .vcf, prior to imputation | 7322630 | See main text |
| **Using filtered .vcf** | | |
| Admixture, PCA | 893961 | r2 > 0.1 on filtered .vcf |
| **Using imputed .vcf** | | |
| GEMMA ULMM | 3021696 | Main imputed .vcf. Allows ≤ 5% missing data, using unmasked reference genome. |
| Genetic relatedness matrix in GEMMA | 344164 | LD-pruned imputed .vcf |
| bgchm | 49019 | See main text |
| AncestryHMM | 44095 | AFD > 0.75 & r2 < 0.2 in 50 bp windows sliding 10 bp |
| Excess heterozygosity analyses | 136590 | All SNPs fixed between parent species |
| **Using all-sites .vcf** | | |
| pixy | 8189072 | See main text |

*Table S4. Differentiation, diversity, and divergence*

| Metric | Quantile | Value | |
|---|---|---|---|
| $d_{xy}$ | 1% | 0.0037 | |
| | 5% | 0.0059 | |
| | 50% | 0.0125 | |
| | 95% | 0.0181 | |
| | 99% | 0.0200 | |
| $F_{ST}$ | 1% | 0.0253 | |
| | 5% | 0.0401 | |
| | 50% | 0.2923 | |
| | 95% | 0.7781 | |
| | 99% | 0.8402 | |
| **Metric** | **Quantile** | ***davidsonii*** | ***newberryi*** |
| π | 1% | 0.0015 | 0.0020 |
| | 5% | 0.0023 | 0.0031 |
| | 50% | 0.0066 | 0.0071 |
| | 95% | 0.0129 | 0.0128 |
| | 99% | 0.0153 | 0.0152 |

*Values represent the 1%, 5%, 50% (median), 95%, and 99% quantiles for $d_{xy}$, $F_{ST}$, and π.*

*Table S5. Floral traits measured in the study and their biological relevance*

| Trait | Biological Relevance |
|---|---|
| Crease to bottom height | Floral mouth geometry; can filter pollinators on size/shape |
| Crease to top height | Floral mouth geometry; can filter pollinators on size/shape |
| Floral crease ratio | Floral mouth geometry; can filter pollinators on size/shape |
| Floral tube length | Restricts access to nectar by tongue/bill length |
| Long stamen length | Sets pollen on pollinator body; mismatch reduces pollen transfer efficiency |
| Long stamen pistil ratio | Herkogamy proxy; affects self-pollen deposistion on stigma and precision of cross-pollen placement |
| Long stamen tube ratio | Determines position of anthers in flower, affecting pollinator contact point |
| Lower petal angle | Landing platform orientation; affects insect approach and positioning during visitation |
| Mouth angle | Influences ease of entry and body orientation |
| Mouth constriction angle | Influences ease of entry and body orientation |
| Mouth height width ratio | Entrance geometry; can filter pollinators on size/shape |
| Nectary area | Nectar production/reward; influences pollinator attraction and visitation rate |
| Pistil length | Receives pollen from pollinator body; mismatch reduces pollen transfer efficiency |
| Pistil tube ratio | Determines position of stigma in flower, affecting pollinator contact point |
| Platform angle | Landing platform orientation; affects insect approach and positioning during visitation |
| Short stamen length | Sets pollen on pollinator body; mismatch reduces pollen transfer efficiency |
| Stamen length ratio | Dimorphic stamen lengths can place pollen on two distinct body parts, affecting pollen transfer |
| Throat inflation ratio | Affects pollinator fit and access to nectar |
| Tube height at base | Affects pollinator fit and access to nectar |
| Tube height at inflection | Affects pollinator fit and access to nectar |
| Tube height at mouth | Affects pollinator fit and access to nectar |
| Tube width at base | Affects pollinator fit and access to nectar |
| Tube width at mouth | Affects pollinator fit and access to nectar |
| Upper petal angle | Influences ease of entry and body orientation |
| Chroma | Color saturation; affects pollinator attraction and discrimination |
| Modified hue | Color; affects pollinator attraction and discrimination |

*Table S6. Random Forest confusion matrix*

| | ***davidsonii*** | ***newberryi*** | **Class Error** |
|---|---|---|---|
| ***davidsonii*** | 1 | 0 | 0 |
| ***newberryi*** | 0 | 1 | 0 |

*Table S7. AIC model comparison*

| Model | df | AIC | ΔAIC |
|---|---|---|---|
| modified_hue ~ F3'5'H | 4 | 1523.385 | -100.313 |
| modified_hue ~ F3'5'H + DFR/ANR | 6 | 1423.072 | 0 |
| modified_hue ~ F3'5'H * DFR/ANR | 8 | 1423.847 | -0.775 |

*Table S8. Genomic clines standard deviations*

| **$SD_c$ (cline center)** | | |
|---|---|---|
| **5%** | **50%** | **95%** |
| 0.5943029 | 0.6315415 | 0.671035 |
| **$SD_v$ (cline gradient)** | | |
| **5%** | **50%** | **95%** |
| 0.1189388 | 0.1289992 | 0.139611 |

*Values represent the 5%, 50% (median), and 95% posterior quantiles of the among-locus variance parameters for genomic cline center ($SD_c$) and slope ($SD_v$).*

*Table S9. Permutation tests for genomic clines*

| Steep clines | | | |
|---|---|---|---|
| **Pseudochromosome** | **Observed** | **Null mean** | **p-value** |
| 1085 | 10 | 112.5 | 1 |
| 1086 | 992 | 1345.3 | 1 |
| 1087 | 333 | 847.5 | 1 |
| 2151 | 4 | 82.7 | 1 |
| 2446 | 1 | 41.6 | 1 |
| 2531 | 2 | 27.6 | 1 |
| 2532 | 667 | 946.3 | 1 |
| 2533 | 1520 | 891.9 | 0* |
| 2684 | 1189 | 581.7 | 0* |
| 2685 | 205 | 416.7 | 1 |
| 2686 | 345 | 590.5 | 1 |
| 2687 | 1649 | 1032.6 | 0 |
| **Shallow clines** | | | |
| **Pseudochromosome** | **Observed** | **Null mean** | **p-value** |
| 1085 | 30 | 69.4 | 1 |
| 1086 | 1745 | 827.0 | 0* |
| 1087 | 404 | 521.9 | 1 |
| 2151 | 127 | 50.9 | 0* |
| 2446 | 10 | 25.5 | 1 |
| 2531 | 8 | 16.8 | 0.996 |
| 2532 | 828 | 579.4 | 0* |
| 2533 | 205 | 548.4 | 1 |
| 2684 | 107 | 357.6 | 1 |
| 2685 | 281 | 255.9 | 0.062 |
| 2686 | 258 | 363.1 | 1 |
| 2687 | 248 | 635.1 | 1 |

*Results are based on 1000 permutations. Observed denotes the number of credibly steep or shallow clines on each pseudochromosome. The null mean is the average genome-wide number of credibly steep or shallow clines, scaled by the proportion of total clines estimated for each pseudochromosome. Statistically significant p-values ($p < 0.05$) are denoted with an asterisk.*

*Table S10. GWAS trait transformations*

| Trait | Transformation | Original-p | Transformed-p |
|---|---|---|---|
| crease_to_bottom_height | sqrt | 3.14E-05 | 8.99E-02 |
| crease_to_top_height | log | 6.28E-10 | 2.81E-03 |
| floral_crease_ratio | log | 9.73E-17 | 4.49E-06 |
| floral_tube_length | log | 1.67E-05 | 1.49E-02 |
| long_stamen_length | none | 3.01E-02 | - |
| longstamen_pistil_ratio | sqrt | 7.29E-03 | 8.55E-03 |
| longstamen_tube_ratio | none | 2.19E-02 | - |
| lower_petal_angle | none | 7.82E-01 | - |
| mouth_angle | none | 5.00E-02 | - |
| mouth_constriction_angle | none | 5.52E-11 | - |
| mouth_height_width_ratio | log | 1.03E-02 | 2.12E-01 |
| nectary_area | sqrt | 1.39E-03 | 4.60E-01 |
| pistil_length | sqrt | 3.59E-02 | 1.08E-01 |
| pistil_tube_ratio | none | 7.68E-01 | - |
| platform_angle | log | 5.01E-05 | 3.38E-01 |
| short_stamen_length | sqrt | 1.74E-02 | 5.52E-02 |
| stamen_length_ratio | log | 8.71E-04 | 2.51E-02 |
| throat_inflation_ratio | log | 3.44E-06 | 8.90E-02 |
| tube_height_at_base | log | 1.53E-10 | 5.78E-05 |
| tube_height_at_inflection | log | 1.44E-11 | 4.07E-06 |
| tube_height_at_mouth | log | 1.61E-10 | 5.84E-05 |
| tube_width_at_base | log | 1.16E-07 | 1.76E-02 |
| tube_width_at_mouth | log | 2.68E-08 | 1.41E-02 |
| upper_petal_angle | none | 2.37E-03 | - |
| chroma | none | 3.40E-01 | - |
| modified_hue | none | 5.66E-10 | - |

*Table S11. GenBank accession information*

| Taxon | GenBank ID | Annotation | Gene |
|---|---|---|---|
| *Arabidopsis thaliana* | NP_178197.1 | cinnamoyl coa reductase | CCR |
| *Arabidopsis thaliana* | KAL9860712.1 | Anthocyanidin reductase | ANR |
| *Arabidopsis thaliana* | NP_199094.1 | dihydroflavonol 4-reductase | DFR |
| *Arabidopsis thaliana* | NP_194455.2 | NAD(P)-binding Rossmann-fold superfamily protein | DFR-like |
| *Buddleja alternifolia* | KAG8383351.1 | hypothetical protein BUALT_Bualt04G0003400 | DFR |
| *Buddleja alternifolia* | KAG8386202.1 | hypothetical protein BUALT_Bualt03G0124500 | DFR-like |
| *Glycine max* | NP_001353986.1 | cinnamoyl-CoA reductase | CCR |
| *Glycine max* | NP_001241913.3 | anthocyanidin reductase 1 | ANR |
| *Glycine max* | NP_001341095.1 | dihydroflavonol-4-reductase | DFR |
| *Glycine max* | XP_003539568.1 | putative anthocyanidin reductase | DFR-like |
| *Orobanche gracilis* | KAL6518447.1 | hypothetical protein OROGR_018949 | DFR |
| *Orobanche gracilis* | KAL6529545.1 | hypothetical protein OROGR_015168 | DFR-like |
| *Orobanche minor* | KAL6583477.1 | hypothetical protein OROMI_005555 | DFR |
| *Orobanche minor* | KAL6553944.1 | hypothetical protein OROMI_019617 | DFR-like |
| *Paulownia fortunei* | KAI3450970.1 | hypothetical protein Pfo_007635 | DFR |
| *Paulownia fortunei* | KAI3457011.1 | hypothetical protein Pfo_013674 | DFR-like |
| *Penstemon barbatus* | AIY51701.1 | dihydroflavonol 4-reductase | DFR |
| *Penstemon barbatus* | Pbar_2022_maker_412243-RA* | Epimerase domain-containing protein | DFR-like |
| *Penstemon davidsonii* | KAK4486022.1 | hypothetical protein RD792_008684 | DFR |
| *Penstemon davidsonii* | KAK4491977.1 | hypothetical protein RD792_002762 | DFR-like |

*Table S11. Cont.*

| | | | |
|---|---|---|---|
| *Penstemon eatonii* | Pe_M00000012379* | Dihydroflavonol 4-reductase | DFR |
| *Penstemon eatonii* | Pe_M00000018217* | Putative anthocyanidin reductase | DFR-like |
| *Penstemon neomexicanus* | AIY51700.1 | dihydroflavonol 4-reductase | DFR |
| *Penstemon smallii* | KAL3818775.1 | hypothetical protein ACJIZ3_004680 | DFR |
| *Penstemon smallii* | KAL3850810.1 | hypothetical protein ACJIZ3_012692 | DFR-like |
| *Perilla frutescens var. hirtella* | KAH6833815.1 | dihydroflavonol 4-reductase | DFR |
| *Perilla frutescens var. hirtella* | KAH6756699.1 | Rossmann-fold superfamily protein | DFR-like |
| *Rehmannia glutinosa* | KAK6125668.1 | hypothetical protein DH2020_040594 | DFR |
| *Rehmannia glutinosa* | KAK6157149.1 | hypothetical protein DH2020_011397 | DFR-like |
| *Sesamum alatum* | KAK4417444.1 | Dihydroflavonol 4-reductase | DFR |
| *Sesamum alatum* | KAK4439852.1 | putative anthocyanidin reductase | DFR-like |

*Information in the Annotation column is the name of the gene in the GenBank database. When unavailable on GenBank (i.e., local blast: marked with asterisk *), we refer to the functional annotation accompanying the genome assembly. The Penstemon eatonii annotations were provided upon request by the authors of Jarvis et al. (2025). The Penstemon barbatus annotations were first published as part of Wessinger et al. (2023) and further modified in Katzer (2024), and are available on NCBI under BioProject PRJNA479669.*

*Table S12. Allele frequency differences at the ANR/DFR candidate gene*

| Scaffold | Position (bp) | AF (davidsonii) | AF (newberryi) | AFD |
|---|---|---|---|---|
| 1086 | 37310824 | 1 | 0 | 1 |
| | 37311543 | 1 | 0.0185185 | 0.9814815 |
| | 37311801 | 1 | 0.0185185 | 0.9814815 |
| | 37311810 | 1 | 0 | 1 |
| | 37311961 | 0.989362 | 0 | 0.989362 |
| | 37313127 | 1 | 0.0185185 | 0.9814815 |
| | 37313131 | 1 | 0.0185185 | 0.9814815 |
| | 37313316 | 1 | 0.0185185 | 0.9814815 |
| | 37313853 | 1 | 0 | 1 |
| | 37313887 | 1 | 0 | 1 |

*Each SNP is a SNP within the ANR/DFR candidate gene with a significant association with floral hue in the GWAS. AF (davidsonii) is the allele frequency of the major allele in P. davidsonii parent samples, AF (newberryi) is that frequency in P. newberryi samples, and AFD is the allele frequency difference calculated between parent populations.*

*Table S13. PCR information*

| Primer Name | Direction | Sequence |
|---|---|---|
| MMS_07 | forward | GTGGCCTCGACTCCTGAATC |
| MMS_08 | reverse | GGTTCCGGCCTATGACTCTG |

*PCR used NEB OneTaq (New England Biolabs), and 35 cycles of the following steps: (1) Denature at 95 degrees for 30 seconds, (2) Anneal at 54 degrees for 30 seconds, (3) Extension at 72 degrees for 60 seconds.*
